# Homoploid Hybrid Speciation in a Marine Pelagic Fish *Megalaspis cordyla* (Carangidae)

**DOI:** 10.1101/2024.08.17.608424

**Authors:** Nozomu Muto, Yong-Chao Su, Harutaka Hata, Nguyen Van Quan, Veera Vilasri, Mazlan Abd. Ghaffar, Ricardo P. Babaran

## Abstract

Homoploid hybrid speciation (HHS) is an enigmatic evolutionary process where new species arise through hybridization of divergent lineages without changes in chromosome number. Although increasingly documented in various taxa and ecosystems, convincing cases of HHS in marine fishes have been lacking. This study presents evidence of HHS in Torpedo scad *Megalaspis cordyla* based on comprehensive genomic, morphological, and ecological analyses. A Principal Component Analysis using thousands of SNPs identified three sympatric clusters in the western Pacific. Genome-wide differentiation between the clusters and the admixed nature of a cluster between the others were evident from population genomic analyses, species tree estimation, mitochondrial DNA divergence, and tests of introgression. Multiple statistical methods for hybrid detection also supported the admixed ancestry of this cluster. Moreover, model-based demographic inference favored a hybrid speciation scenario over introgression. Examination of occurrence data and ecologically relevant morphological characters suggested ecological differences between the clusters, potentially contributing to reproductive isolation and niche partitioning in sympatry. The clusters are morphologically distinguishable and thus can be taxonomically recognized as separate species. The hybrid cluster is restricted to the coasts of Taiwan and Japan, where all three clusters coexist. The parental clusters are additionally found in lower latitudes such as the coasts of the Philippines, Vietnam, Thailand, and Malaysia, where they display non-overlapping distributions. Given the geographical distributions, estimated times of the species formation, and patterns of historical demographic changes, we propose that the Pleistocene glacial cycles were the primary driver of HHS in this system. Based on this argument, we develop an ecogeographic model of HHS in marine coastal ecosystems, including a novel hypothesis to explain the initial stages of HHS.

---

Hybridization can drive adaptation and species diversification. One possible outcome of hybridization is hybrid speciation, where interbreeding between diverged species results in a new population evolving independently from the parental species (Coyne and Orr 2004; Mallet 2007; Peñalba et al. 2024). In allopolyploid hybrid speciation, a hybrid population arises with whole-genome duplication, reported frequently in plants (Wood et al. 2009). In contrast, homoploid hybrid speciation (HHS) occurs without changes in ploidy, with the nascent hybrid population possessing a recombinant genome of parental species (Mallet 2007). There is much debate about the criteria for a case to be regarded as HHS. An emerging view is that at least the first and second of the following criteria must be satisfied, whereas a more stringent definition requires the third as well: 1) the species arose via hybridization with detectable genomic signatures, 2) intrinsic or extrinsic reproductive isolation from parental species has been established, and 3) the reproductive barriers directly resulting from hybridization (Abbott et al. 2013; Schumer et al. 2014; Nieto Feliner et al. 2017; Long and Rieseberg 2024). Cases of HHS fulfilling the first two criteria are increasingly documented. However, it is still rare compared to other hybridization-related phenomena, such as the formation of hybrid zones and introgression. Moreover, reported cases are strongly biased towards terrestrial and freshwater organisms (Abbott et al. 2013; Yakimowski and Rieseberg 2014; Taylor and Larson 2019 and references therein).

A major challenge in HHS is the persistence of the nascent hybrid population. Immediately after formation, it faces ecological competition with parental species and the swamping effects of gene flow (Buerkle et al. 2000). The hybrid population can overcome both and persist on an evolutionary timescale by acquiring a novel niche not occupied by the parental species. The acquisition of a novel niche drives HHS in several ways. First, it may directly cause extrinsic pre- or postzygotic reproductive isolation between the hybrid and parental species (Rieseberg et al. 2003; Schwarz et al. 2005). Second, it enables the hybrid and parental species to coexist in sympatry by alleviating competition through niche partitioning (Lamichhaney et al. 2018; Olave et al. 2022). Third, it may allow the hybrid species to occupy a spatially separated habitat from the parental species, facilitating allopatric or parapatric divergence (Donovan et al. 2010; Brennan et al. 2012; Lopes et al. 2023). Supporting the importance of the third role, the independence of the hybrid species in most compelling cases of homoploid hybrid speciation seems to rely on geographical separation from at least one parental species (Rieseberg et al. 2003; Mavárez and Linares 2008; Larsen et al. 2010; Brennan et al. 2012; Kang et al. 2013; Lamichhaney et al. 2018; Maier et al. 2019; Noguerales and Ortego 2022; but see Olave et al. 2022 and Rosser et al. 2024 for counterexamples). Theory suggests that the recombinant hybrid genome can express transgressive phenotypes via additive and/or epistatic effects of parental genetic variants, leading to the exploitation of novel niches (Rieseberg et al. 2003; Gross and Rieseberg 2005; Mallet 2007; Abbott et al. 2013). However, in what ecogeographic settings the niche divergence leads to reproductive or geographical isolation remains largely unknown.

The Pleistocene glacial cycles significantly impacted the evolutionary histories of organisms at and below the species level across various ecosystems and taxa. In terrestrial ecosystems, ice sheets formed in high altitudes and latitudes during glacial periods disrupted existing habitats, while glacial retreat during interglacial periods created new, unoccupied habitats. Coupled with temperature changes, these habitat dynamics resulted in allopatric divergence, secondary contact, altitudinal and latitudinal range shifts, population size changes, and local adaptation (Hewitt 2004). Mountain systems are among the most well-studied (Hewitt 2000, 2004), including a few possible cases of homoploid hybrid speciation (Gompert et al. 2006; Maier et al. 2019). Marine coastal areas are another ecosystem strongly influenced by historical climate changes. Eustatic sea level changes cyclically disrupted and opened habitats for marine organisms in shallow waters, creating barriers or corridors to gene flow (Bowen et al. 2016). Changes in water temperature and associated environmental factors caused latitudinal range shifts, population size changes, and adaptive responses, similar to those ongoing under contemporary climate change (Kiessling et al. 2012; Neiva et al. 2014; Donelson et al. 2019). These, in turn, resulted in various hybridization-related phenomena in coastal organisms via secondary contact: introgression (Duranton et al. 2020), formation of stable hybrid zones (Hirase et al. 2021), and divergence with gene flow (Muto and Kai 2023). Nonetheless, HHS in this environment has not been documented to date.

Here, we provide genomic and phenotypic evidence for HHS in the torpedo scad *Megalaspis cordyla* (Linnaeus, 1758) in the western Pacific, representing the first compelling case of this phenomenon in a marine pelagic fish. The torpedo scad is a pelagic marine predator in the family Carangidae (Teleostei), usually reaching 40 cm in total length. Widely distributed in tropical to temperate coastal waters throughout the Indian Ocean and the western Pacific, it is an important fisheries resource in the surrounding countries (Smith-Vaniz 1999). Previous phylogeographic studies based on mitochondrial DNA (mtDNA) sequences suggested historical vicariance followed by secondary contact in the western Pacific region. Specimens from the southwestern part of the region (the Pacific coast of the Asian continent, from Vietnam to Malaysia) and the southeastern part (the islands of the Philippines and Indonesia) represent reciprocally monophyletic lineages, whereas specimens from the northern part (Taiwan) represent both lineages (Su et al. 2020; Muto et al. 2021). In the present study, we provided evidence for genome-wide divergence between the two lineages. We also discovered a third genetic cluster restricted to locations where the two lineages are sympatric. We tested the hypotheses that this third cluster originated from the admixture of the other two and that they represent separate species, using detailed genomic, ecological, and morphological analyses. Based on the results, we suggest that the Pleistocene glacial cycles primarily drove HHS in the present system. Furthermore, we develop an ecogeographic model of HHS in coastal marine environments, where demographic dynamics under climate change and adaptation across latitudinal environmental gradients play critical roles. This model bridges a major gap in the understanding of how the initial stages of HHS proceed.

## MATERIALS AND METHODS

A full description of methods is provided in Online Appendix 1.

### Specimens

We analyzed a total of 160 specimens of *M. cordyla* collected from various localities in the western Pacific (Fig. 1a), where two distinct mtDNA lineages have been discovered (Su et al. 2020; Muto et al. 2021). For genotyping, we used tissues stored in natural history collections in respective countries (Table S1). Total genomic DNA was extracted from the fin or muscle tissue of each specimen using the Favorgen Tissue Genomic DNA Extraction Mini Kit (Favorgen). The specimens from Vietnam, Thailand, Malaysia, and the Philippines were analyzed in the previous study (Muto et al. 2021).

**Fig. 1.**
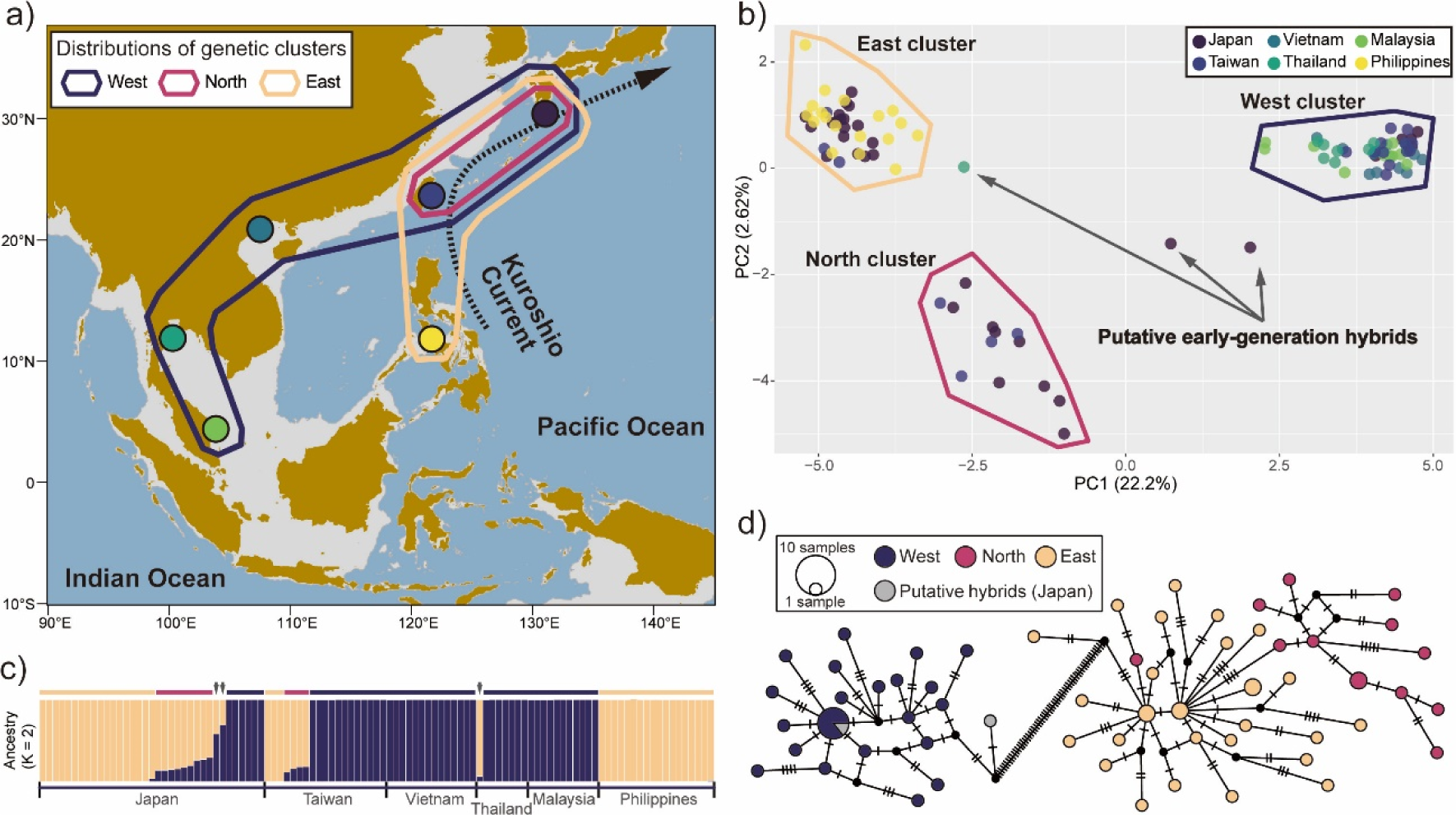
a) Sampling sites (circles) and distributions of three genetic clusters (polygons) of *Megalaspis cordyla* in the western Pacific. The grey-shaded area represents regions currently shallower than 130 m, repeatedly exposed during Pleistocene glacial cycles. The dotted arrow indicates the approximate route of the Kuroshio Current. b) Scatter plots for the principal component analysis (PCA) based on the genome-wide SNPs, with polygons and arrows indicating the three genetic clusters and their putative early-generation hybrids, respectively. The clusters are named according to their geographical distributions. c) Result of the STRUCTURE analysis assuming the number of clusters (*K*) = 2, which was the optimal according to the delta *K* values. Horizontal bars and arrows indicate individual assignments to the three clusters and putative early-generation hybrids, respectively, based on the PCA. The color code for the horizontal bars matches that for the PCA plot polygons. Note that individuals in the North cluster and putative early-generation hybrids show admixed ancestry. d) Statistically parsimonious network of haplotypes of the concatenated control region and cytochrome b gene sequences of the mitochondrial genome. Circle/pie-chart size is proportional to haplotype frequency. Black dots and a hatch mark along branches represent unobserved hypothetical haplotypes and one mutational step, respectively.

### Mitochondrial DNA Genotyping

We obtained partial sequences of the mitochondrial cytochrome b gene (CYTB: 873 bp) from all specimens. Additionally, we obtained partial sequences of the following seven loci from a subset of the specimens: cytochrome c oxidase subunits 1 (COI: 612 bp) and 2 (COII: 657 bp), NADH dehydrogenase subunits 1 (ND1: 828 bp), 2 (ND2: 531 bp), 4 (ND4: 828 bp), and 5 (ND5: 801 bp), and the control region (CR: 760 bp) (Table S1). We amplified the COI and CYTB sequences following Ivanova et al. (2007) and Cárdenas et al. (2005), respectively. The remaining loci were amplified using newly developed primers (Table S2). We edited and aligned the sequences using BioEdit v7.0.5.3 (Hall 1999) and CLUSTAL X (Larkin et al. 2007).

### Genome-Wide SNPs Genotyping

We genotyped 115 specimens using MIG-seq (Suyama and Matsuki 2015; Suyama et al. 2022), a reduced representation sequencing method targeting SNPs in the inter-simple sequence repeat region (ISSR). We performed the first PCR using 12 multiplex primers to amplify various ISSR across the genome (Suyama and Matsuki 2015; Suyama et al. 2022). The first PCR used 12 multiplex primers to amplify various ISSRs across the genome. The second PCR added Illumina sequencing adapters and a unique dual index. We pooled the second PCR products, size-selected for 400–800 bp, purified, quality-checked, quantified, and sequenced on an Illumina Novaseq 6000 in 150 bp paired-end mode at Novogen Co. Ltd., China, yielding 148M reads.

We processed the raw MIG-seq reads by the single-end mode of fastp v0.20.1 (Chen et al. 2018), merging the processed reads 1 and 2 into a single FASTQ file, and mapped them to the *Caranx ignobilis* reference genome (GenBank JAFHLA000000000.1) (Pickett et al. 2022) using BWA-mem v0.7.17 (Li 2013) with default settings. Unmapped or low-quality reads were removed using Samtools v1.12 (Danecek et al. 2021). The resulting BAM files were used to prepare 5 datasets for downstream analyses (see the next paragraph). We also downloaded the whole-genome shotgun reads of *C. ignobilis* (a different individual from the reference) and *Caranx melampygus* from the NCBI Short Read Archive (accession nos.SRR1002875 and SRR1002876) (Santos 2014) as outgroups. We processed these reads using the paired-end mode of fastp, mapped them to the reference genome using BWA-mem with default settings, and removed unmapped, low-quality, or duplicate reads using Samtools. The resulting BAM files were used for the analyses requiring outgroups (datasets 2– 5: see below).

The following 5 datasets were prepared for downstream analyses: 1) 2,926 unlinked biallelic SNPs in all available (105) specimens of *M. cordyla* without an outgroup for population genetic analyses using various tools and ancestry estimation using HIest v2.0 (Fitzpatrick 2012); 2) 6,789 unlinked biallelic SNPs in 24 specimens (8 per genetic cluster) with *C. ignobilis* as an outgroup for admixture detection using HyDe v1.0.0 (Blischak et al. 2018) and ADMIXTOOLS v7.0.2 (Patterson et al. 2012); 3) 3,179 unlinked biallelic SNPs in 9 specimens (3 per genetic cluster) with *C. ignobilis* as an outgroup for divergence time estimation using SNAPP v1.5.0 (Bryant et al. 2012); 4) 6,341 biallelic SNPs in the same 24 specimens, polarized using the *C. ignobilis* and *C. melampygus* genomes as outgroups, for demographic model selection using fastsimcoal v2.7.0.9 (Excoffier et al. 2013, 2021); and 5) 22,326 biallelic SNPs (linked and unlinked) and 920,184 monomorphic sites in the same 24 specimens, polarized using the *C. ignobilis* and *C. melampygus* genomes as outgroups, for demographic parameter estimation using fastsimcoal 2 and historical population size estimation using Stairway Plot v2.1.2 (Liu and Fu 2020). Note that monomorphic sites and linked SNPs in the 5th dataset are needed for direct parameter estimations using fastsimcoal 2, whereas only unlinked SNPs are used for the model selection (Excoffier et al. 2013). For each dataset, we called SNPs using Stacks v2.5.4 (Catchen et al. 2013) or *mpileup* implemented in bcftools v1.8 (Danecek et al. 2021). We then filtered the SNPs and created final vcf files using VCFtools v0.1.16 (Danecek et al. 2011) and R package ‘SNPfiltR’ v1.0.0 (DeRaad 2022) on R v4.2.2 (R Core Team 2022).

We used *easySFS.py* (https://github.com/isaacovercast/easySFS) to calculate the unfolded site frequency spectrum (SFS) from the 4th and 5th datasets, assuming the reference alleles fixed in both outgroups represent the ancestral state. The resulting one-dimensional SFS from the 5th dataset was used for the historical population size estimation by Stairway Plot 2, whereas multi-dimensional SFS was used for the demographic inference by fastsimcoal 2. For input file preparations for the other analyses, we used PGDSpider v2.1.1.2 (Lischer and Excoffier 2012) to convert the vcf files to appropriate formats.

### Population Genetic Analyses

We constructed minimum-spanning haplotype networks of mtDNA sequences using PopART v1.7 (Leigh and Bryant 2015).

The following population genetic analyses were conducted using 1st dataset of the genome-wide SNPs. We performed a principal component analysis (PCA) using the ‘adegenet’ package v2.1.4 in R (Jombart 2008). We used STRUCTURE v2.3.4 (Pritchard et al. 2000) to identify admixed individuals with 10 replicate runs for each predefined number of genetic clusters (*K*) = 1–8, each with 100,000 Markov Chain Monte Carlo (MCMC) generations after 100,000 generations of burn-in. These runs were parallelized using StrAuto v1.0 (Chhatre and Emerson 2017). Statistical support for *K* values was evaluated by calculating the Δ*K* value (Evanno et al. 2005) using Structure Harvester v0.6.94 (Earl and vonHoldt 2012). The results were summarized and visualized with ‘pophelper’ v2.3.1 in R (Francis 2017). We used SplitsTree v5.3.0 (Huson and Bryant 2006) to build an unrooted phylogenetic network based on the Neighbor-Net algorithm.

The population genetic analyses showed one genetic cluster was genetically intermediate to the other two clusters (Fig. 1b, c). This intermediate cluster is hereafter called the “North” cluster, while the others are the “East” and “West” clusters, according to their geographical distributions (Fig. 1a). Putative early-generation hybrids (F1, F2, or backcross) between these clusters were also detected. Excluding these putative hybrids, we calculated locus-specific pairwise *F_ST_* between the clusters using *populations* implemented in Stacks, and pairwise *F_ST_* across all loci and associated *p*-values using Arlequin v3.5.2.2 (Excoffier and Lischer 2010).

We estimated historical population size changes in each cluster using Stairway Plot v2.1.2 (Liu and Fu 2020) based on the 5th dataset. We assumed a mutation rate of 1E^−8^ per site per generation (Tine et al. 2014) and a generation time of 1.2 years (Froese and Pauly 2023). Other settings were default or those recommended by the author of the software.

### Divergence Time Estimation

We estimated the divergence times of the three genetic clusters using genome-wide SNP dataset 3 and mtDNA sequence data. The former was done under the multispecies coalescent model using the SNAPP v1.5.0 package (Bryant et al. 2012) of BEAST v2.7.1 (Bouckaert et al. 2014), following Stange et al. (2018). Time calibration was based on the estimated node age of 6.349 (95% CI: 4.434-8.698) million years ago (Ma) for the most recent common ancestor (MRCA) of *C. ignobilis* and *M. cordyla* (Glass et al. 2023), approximated by a lognormal distribution with a mean of 6.349 Ma, a standard deviation of 0.205, and an offset of −0.01. We performed two independent runs of 50,000,000 MCMC generations, sampling every 10,000 steps and discarding the first 10% of the trees as burn-in, then combined the posterior tree distributions from the two runs. We estimated a maximum clade credibility tree using TreeAnnotator v2.7.1 (Bouckaert et al. 2014) and visualized the tree using FigTree v1.4.4 (http://tree.bio.ed.ac.uk/software/figtree/). The estimation based on the concatenated sequences of the 8 mtDNA loci was done using BEAST v2.7.1. We partitioned the sequences by locus and estimated the substitution model for each partition using the bModelTest v1.3.3 (Bouckaert and Drummond 2017), which averages over different substitution models in BEAST. We assumed the optimized relaxed clock model and the Yule model for the clock model and the tree prior, respectively. We used the mitogenome sequence of *C. ignobilis* (accession No.: NC_022932) as an outgroup. Time calibration, MCMC simulations, convergence check, tree estimation, and visualization were the same as described above.

### Ancestry Estimation and Admixture Detection

We estimated the ancestries of the individuals in the North cluster and the putative early-generation hybrids using the ‘HIest’ package v2.0 (Fitzpatrick 2012) in R. This program estimates the ancestries of target individuals relative to predefined parents via log-likelihood maximization, using target genotypes and parental allele frequencies. First, we assumed the East and West clusters as potential parents and used 108 unlinked SNPs with allele frequencies differing by >0.5 between them. Second, we estimated the ancestries of putative hybrids only, assuming the three clusters as potential parents, using 130 SNPs with allele frequencies differing by >0.5 between one or more pairs of the clusters. In the first analysis, we estimated the ancestry index (*S*) and the inter-class heterozygosity (*H*) using the *HIest* function, employing the SANN method for searching the likelihood surface, 500 MCMC generations, and the option surf = TRUE. *S* represents the proportion of an individual’s alleles derived from one predefined parental population (here, the East cluster), equivalent to the hybrid index. *H* indicates the proportion of loci with one allele from each parental population. Expected *H* values vary between classes of early-generation hybrids (F1: 1.0; F2 and backcross: 0.5), decreasing if succeeding reproduction occurs exclusively among hybrids and alleles are lost by drift. We also estimated the likelihoods for six possible genotype classes resulting from 2 generations of inter- or intraspecific crosses (parental, F1, F2, and backcrosses) using the *HIclass* function. We compared the maximum likelihood of the six classes with that of the maximum likelihood estimation of *S* and *H* using the *HItest* function. For the second analysis, we estimated the ancestry indices *S*_East_, *S*_West_, and *S*_North_ of the putative early-generation hybrids using the *threeway* function. These indices are equivalent to *S*, representing the proportion of alleles derived from the three predefined parental populations. We also calculated the likelihoods for 15 possible genotype classes in the first two generations of hybridization between pairs of parental populations using the *thirdclass* function and compared the maximum likelihood with that obtained from the *threeway* analysis using the *HItest3* function.

We tested if the North cluster originated from an admixture of the East and West clusters by three methods based on the 3rd dataset of the genome-wide SNPs. First, we used HyDe v1.0.0 (Blischak et al. 2018), which assesses the evidence for hybridization under a model of hybrid speciation and estimates the associated ancestry proportions in a putative hybrid population from two parental populations (г and 1−г). We tested for hybridization at the population level for each cluster, leaving the remaining clusters as putative parental populations, using the *run_hyde.py* script. We then performed 500 bootstrap resampling using the *bootstrap_hyde.py* script to detect individual variation in г within a hybrid population. Second, we calculated the *f*_3_ statistic using ‘admixr’ v0.9.1 (Petr et al. 2019). The *f*_3_ statistic is a three-population statistic based on branch lengths between populations. A significantly negative *f*_3_ statistic indicates that a predefined outgroup population is a mixture of two ingroup populations (Patterson et al. 2012). We calculated the *f*_3_ statistics designating each cluster as an outgroup. Third, we calculated the *D*-statistic using ‘admixr’ to test for introgression between the clusters. The *D*-statistic detects introgression between non-sister taxa under the user-defined topology by asking if the allele-sharing pattern between the taxa deviates from what is expected under incomplete lineage sorting without introgression (Patterson et al. 2012). Given the possible hybrid origin of the North cluster (see Results), two strictly bifurcating topologies were possible for the user-defined topology, i.e., one with a sister relationship between the East and North clusters, i.e., (East,North),West), and another between the West and North clusters, (West,North),East). These were referred to as topologies 1 and 2, respectively. We calculated the *D*-statistics for both.

### Demographic Model Selection and Parameter Estimation

We inferred the demographic history of *M. cordyla* by testing alternative scenarios using fastsimcoal v2.7.0.9 (Excoffier et al. 2021). Given the admixed nature of the North cluster, we aimed to determine if it originated from an admixture of the East and West clusters (“hybrid speciation” scenario) or an instantaneous introgression event following divergence (“pulse introgression”). We also considered a null scenario of strict bifurcation between the clusters without a pulse introgression event (“bifurcation”), following Noguerales and Ortego (2022). Alternative topologies 1 and 2, i.e., (East,North),West) and (West,North),East), were assumed for the pulse introgression and bifurcation scenarios, resulting in five scenarios. These scenarios were simulated with or without gene flow between the clusters before (ancestral gene flow) and after (contemporary gene flow) the formation of the North cluster, resulting in 20 models (Fig. S1).

We performed 100 independent fastsimcoal runs for each demographic model using the 4th dataset for model selection. Each run comprised 100 expectation-conditional maximization (ECM) cycles with 100,000 coalescent simulations. The run with the smallest difference between the maximum expected likelihood and the maximum observed likelihood was considered the best fit. For model selection, we compared 1) Akaike’s information criterion (AIC) based on the best-fit run of each model, and 2) the likelihood distributions from 100 expected SFS, generated by 1,000,000 coalescent simulations under the parameters estimated from the best-fit run (Meier et al. 2017). Under the selected model, we then estimated demographic parameters using the 5th dataset. To obtain 95% confidence intervals (CIs) for each parameter, we generated 100 parametric bootstrap replicates of SFS under the maximum likelihood parameters of the best-fit run following the program manual. The parameters were re-estimated using 100 independent runs for each bootstrapped SFS with the same settings, except that the initial values were set to the maximum likelihood parameters of the best-fit run, and the number of ECM cycles was set to 50. We rescaled the estimated parameters assuming a mutation rate of 1E^−8^ per site per generation and a generation time of 1.2 years (see above).

### Morphology

We examined 37 linear measurements and 17 countable characters in 37 specimens from Japan, Taiwan, Vietnam, and Thailand (Tables S1, S3). This analysis aimed to determine if the genetic clusters 1) differ in ecologically relevant morphological characters, suggesting ecological divergence, and 2) can be recognized as separate species in a taxonomic sense. Linear measurements were made to the nearest 0.01 mm using a digital vernier caliper. Methods for measuring and counting generally followed Marr and Schaefer (1949), Gibbs and Collette (1967), and Muto et al. (2016). Linear measurements are expressed as % of the Standard length (SL). To illustrate the variation in overall morphology, we performed a PCA of 24 characters with no missing data using the R package ‘FactoMineR’ v2.11 (Lê et al. 2008). All linear measurements were corrected for allometry using the R package ‘GroupStruct’ v0.1.0 (Chan and Grismer 2022) before performing the PCA.

### Ecology

To investigate potential ecological divergence between the genetic clusters, we re-analyzed the ecological data of *M. cordyla* in Taiwan reported by Su et al. (2020). The data include sampling season and location, individual body length and weight, gonad weight, gonad somatic index (GSI: [gonad weight/body weight] × 100), and sex (Table S4), along with the mtCR sequences of the specimens. Su et al. (2020) compared the ecological factors between two major lineages of mtCR sequences. In the present study, we show that one of these two major lineages corresponded to the West cluster, while the other corresponded to the East and North clusters (see Results). Additionally, we demonstrate that the East and North clusters can be distinguished by a minor mtCR divergence within the major lineage. These observations allowed us to reclassify the specimens examined by Su et al. (2020) into the three clusters based solely on the mtCR sequences reported by the authors, albeit with some uncertainty. First, we tested the association between the ecological factors and genetic clusters under the Random Forest model using the R package ‘ranger’ v0.16.0 (Wright and Ziegler 2017). We assessed the importance of each ecological factor using the mean decrease in accuracy and estimated the significance of this importance metric with a permutation test using the R package ‘rfPermute’ v2.5.2 (Archer 2021). Second, we performed a hierarchical cluster analysis to create a dendrogram of the specimens based on the ecological data. We calculated the pairwise Gower’s distance between each specimen and constructed Ward’s hierarchical agglomerative clustering using the R package ‘stats’ v4.2.2 (R Core Team 2022). Third, we performed a Factor Analysis of Mixed Data (FAMD) using the R package ‘FactomineR’.

## RESULTS

### Population Genetic Analyses

The first two PCA axes separated the specimens into three distinct clusters: 1) individuals from Japan to Malaysia along the east coast of the Eurasian continent, 2) individuals from Japan to the Philippines along the Kuroshio Current, and 3) individuals from Japan and Taiwan (Fig. 1a, b). These clusters are referred to as the West, East, and North clusters, respectively, based on their geographical distributions. The East and West clusters occupied opposite positions along PC 1, with the North cluster in an intermediate position. Principal component 2 further separated the North cluster from the others. Two individuals from Japan were intermediate between the North and West clusters and are hereinafter referred to as putative early-generation hybrids. One individual from Thailand was plotted near the East cluster, while all other individuals from Thailand were included in the West cluster. This individual from Thailand is also regarded as a putative early-generation hybrid. In the STRUCTURE analysis, the most likely number of *K* was 2 (Fig. S2). Assuming *K* = 2, individuals of the North cluster showed signatures of admixture between the East and West clusters, with a greater ancestry contribution from the former (Fig. 1c). The putative early-generation hybrids from Japan showed intermediate ancestry between the North and West clusters, while that from Thailand was intermediate between the North and East clusters. Assuming *K* = 3–8, individuals in the North cluster showed admixed ancestry from three clusters: two corresponding to the East and West clusters, and an additional minor cluster (Fig. S2). The Neighbor-Net network showed similar patterns regarding the separation of the East, West, and North clusters, the intermediate nature of the North cluster, and the positions of the putative early-generation hybrids (Fig. S3). Pairwise *F_ST_* across all SNPs was greatest in the East-West comparison, followed by the West-North and East-North comparisons (0.2989, 0.2167, and 0.0861, respectively); all comparisons were significant (Table S5). Significant locus-by-locus pairwise *F_ST_* was most frequently observed in the East-West comparison, followed by the West-North and East-North comparisons (Fig. S4). Several SNPs showed significant differentiation in both East-North and West-North comparisons, highlighting the unique evolutionary history of the North cluster at these loci. In terms of allele frequency, a total of 130 SNPs were highly differentiated (i.e., differences >0.5) between one or more cluster pairs (Fig. S5; Table S6). The majority of these SNPs (95 loci) were highly differentiated in the East-West comparison, with the North cluster showing intermediate allele frequencies. For 56 of these 95 SNPs, the North cluster’s allele frequency was closer to the East cluster, while for others, it was closer to the West cluster, suggesting that alleles from the East and West clusters were alternatively sorted in the North cluster following admixture. In contrast, the North cluster was highly differentiated from both the East and West clusters at 4 SNPs, due to the alleles reaching high frequency in the North cluster while being almost absent in the East and West (0.56–0.67 in the North vs. 0–0.05 in East and West).

The population sizes of the East and West clusters were inferred to have declined approximately 70–100 thousand years ago (Ka), while the North cluster remained stable (Fig. S6). Subsequently, all three clusters expanded in population size towards the present time.

### mtDNA Differentiation

The minimum spanning network based on concatenated sequences of CYTB and CR (873 + 760 bp) consisted of three major haplogroups, each corresponding to one of the three genetic clusters (Fig. 1d). We refer to these haplogroups as the East, West, and North haplogroups, corresponding to their respective clusters. The East-West and East-North divergences of the haplogroups were equivalent to 58 and 4 mutations, respectively. However, the East and North clusters were not reciprocally exclusive: one individual of the North cluster was included in the East haplogroup. Moreover, the East-North divergence disappeared when only the CYTB sequences or concatenated sequences of the seven protein-coding genes were analyzed (Fig. S7), indicating the divergence is due to variation in CR sequences. The putative early-generation hybrids from Japan were included in the West haplogroup (Fig. 1d). We were unable to obtain the CR sequence for the putative hybrid from Thailand; this individual was included in the “East + North” haplogroup in the CYTB-only network (Fig. S7).

### Ancestry Estimation and Admixture Detection

In the *HIest* analysis, assuming the East and West clusters as potential parents, *S* and *H* for individuals in the North cluster were 0.51–0.76 and 0.10– 0.35, respectively (Fig. 2a). The values of *S* suggest a greater ancestry contribution from the East cluster, while those of *H* suggest that the individuals are later-generation hybrids (>F_2_), having exclusively reproduced within the same cluster. *HIclass* assigned each individual in the North cluster to either backcross to the East cluster or F_2_, but the likelihood ratio test by *HItest* rejected these simple assignments in favor of the more nuanced results of *HIest* (Table S7). Compared to the North cluster individuals, the putative early-generation hybrids from Japan showed lower *S* (i.e., more closely related to the West cluster) and higher *H*, whereas the hybrid from Thailand showed greater *S* and lower *H*. *HIclass* assigned them to F_2_ and backcross to the East cluster, respectively, but *HItest* again rejected these assignments (Fig. 2a). In the *threeway* analysis, where the three clusters were assumed as potential parents, the putative early hybrids from Japan and Thailand showed mixed ancestries predominantly between the North and West clusters and between the North and East clusters, respectively (Table S8). *Thirdclass* assigned them to F_2_ between the North and West clusters and F_2_ between the North and East clusters, respectively; *HItest3* did not reject these assignments.

**Fig. 2.**
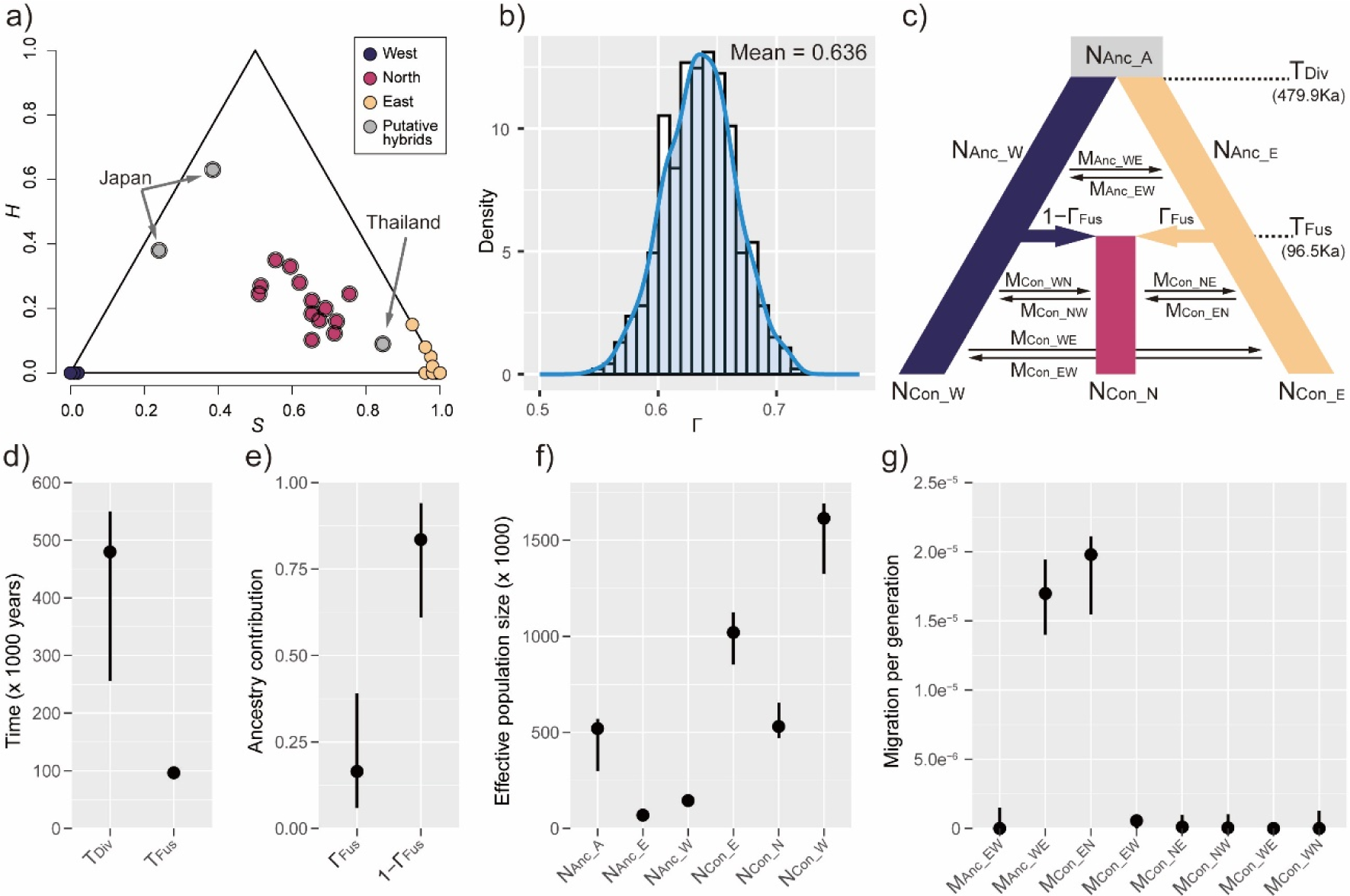
a) Estimates of ancestry index (*S*), equivalent to hybrid index, and inter-class heterozygosity (*H*) by HIest, assuming the East and West clusters as potential parents. Individual assignments to the three clusters and putative early-generation hybrids are based on the PCA (Fig. 1b). Double circles indicate specimens for which simple classification as one of 6 genotype classes within 2 generations of inter- or intraspecific crosses (parentals, F1, F2, and backcrosses) was rejected by *HItest*. See Table S8 for results assuming the three clusters as potential parents. b) Density plot of estimated gamma (Γ) values for the North cluster across 500 individual bootstraps in HyDe. Γ and 1-Γ represent estimated ancestry contributions from the East and West clusters, respectively. c) A schematic diagram of the best-fit demographic model, i.e., hybrid speciation with ancestral and contemporary gene flow between clusters, tested using Fastsimcoal 2. Parameters include; N: ancestral (Anc) or contemporary (Con) effective population size of the common ancestor (_A) or the three genetic clusters (_E / _W / _N); T: time of divergence (Div) or fusion (Fus) of the East and West clusters; Γ: ancestry contribution from the East (ΓFus) or West (1−ΓFus) cluster upon fusion; M: ancestral (Anc) or contemporary (Con) gene flow between clusters, with the letters following the underscore indicating the direction of the gene flow (e.g., M_Con_EW_ represents contemporary gene flow from the East to West cluster forward in time). See Fig. S1 for all 20 models tested and Fig. S9 and Table S10 for model evaluation by likelihood and Akaike’s Information Criterion values. d)–g) Demographic parameters estimated using Fastsimcoal 2 under the best-fit model. Dots and vertical bars represent maximum likelihood point estimates and 95% confidence intervals based on 100 bootstrap replicates. Timings of demographic events are in units of years. Effective population sizes are in units of the number of alleles. Migration rates are in units of proportions of alleles moving per generation. See Table S11 for tabulated values.

Admixture detection using HyDe was significant only for the North cluster (*Z*-score = 2.584, *p* = 0.0049; Table S9), supporting its hybrid origin. Bootstrap resampling of the individuals in the North cluster showed an unimodal distribution of г values, indicating uniform admixture across individuals (Fig. 2b). This uniform admixture is consistent with hybrid speciation rather than ongoing introgression (Blischak et al. 2018). The departure of the г values from 0.5 (mean = 0.636; 95% CI = 0.633–0.638) suggested a greater ancestry contribution from the East cluster. The *f*_3_ statistic was negative when the North cluster was designated as an outgroup, supporting its admixed ancestry (*f*_3_ = - 0.023, *Z*-score = −4.338; Table S9). Finally, the *D*-statistics detected gene flow between both West-North and East-North combinations (West-North: *D* = 0.185, *Z*-score = 7.079; East-North: *D* = 0.296, *Z*-score = 11.389; Table S9).

### Divergence Time Estimation

The East and North clusters formed a single clade, leaving the West cluster in another clade, in both the species tree based on genome-wide SNPs and the mtDNA phylogeny (Fig. S8). The MRCA age of the three clusters was estimated to be 0.269 Ma (95% CI: 0.168–0.387) based on genome-wide SNPs and 1.333 Myr (0.244–3.208) based on mtDNA phylogeny. The divergence between the North and East clusters was estimated at 0.126 Ma (0.075–0.184) and 0.196 Ma (0.015–0.574), respectively. All estimated divergence ages fell within the Pleistocene. Note that the East and North clusters were not reciprocally monophyletic in the mtDNA phylogeny; the divergence time mentioned above is the node age for two clades approximately corresponding to the East and North clusters.

### Demographic Model Selection and Parameter Estimation

The hybrid speciation model with gene flow between clusters both before and after the formation of the North cluster (i.e., ancestral and contemporary gene flow) was the best fit among the 20 models tested, based on the highest log-likelihood and lowest AIC (Fig. 2c; Fig. S9; Table S10). The other three hybrid speciation models, differing in ancestral and contemporary gene flow, were not better supported than the pulse-introgression models assuming a sister relationship between the West and North clusters (topology 2) (Fig. S9; Table S10). The models assuming topology 2 were favored over the alternative topology 1 if accompanied by a pulse introgression event or contemporary gene flow.

Under the best-fit model, the divergence and fusion of the East and West clusters were estimated to have occurred 479.9 Ka (95% CI: 256.1–549.9) and 96.5 Ka (94.9–106.4), respectively (Fig. 2d; Table S11), both within the Pleistocene. The ancestry contribution from the East cluster (Γ_Fus_) upon the fusion was less than that from the West cluster (1−Γ_Fus_) (Fig. 2e). Effective population size was greater in the West than in the East cluster, both before and after the fusion, with both clusters expanding over time (Fig. 2f). Contemporary population size was smallest in the North cluster, consistent with its restricted geographical range. Gene flow between the clusters was highly asymmetric; contemporary gene flow from the East to North cluster was orders of magnitude greater than that in the opposite direction and other cluster combinations (Fig. 2g; Table S11). Similarly, ancestral gene flow from the West to East cluster was significantly greater than that in the opposite direction.

### Ecology

A NJ tree based on the 592 bp alignment of the mtCR sequences, including both the present and Su et al’s. (2020) sequences, showed that 20, 55, and 15 haplotypes reported by Su et al. (2020) are most likely identified as the East, West, and North clusters, respectively (Fig.S10; Table S4). Based on this identification, we tested possible ecological divergence between the three clusters. The Random Forest model showed that sampling location and season were significant predictors of the cluster (*p* < 0.05) (Fig. 3a), suggesting that the clusters tend to be found in different locations and seasons along the coast of Taiwan. This implies habitat segregation or differences in migration patterns contribute to reproductive isolation between the clusters. The Random Forest proximity scores of the three clusters, reduced to two dimensions through multidimensional scaling, largely overlapped (Fig. 3b), whereas the individual assignments to the clusters predicted from the Random Forest model were more accurate for the North cluster than for the other two clusters (Fig. 3c). This suggests that the ecological divergence is more pronounced between the North and the other clusters. In contrast, the dendrogram constructed through the hierarchical cluster analysis could not fully explain the split of expected East, West, and North clusters (Fig. S11a). Similarly, the three genetic clusters were not clearly separated on the first four dimensions (explaining 61.5% of the total variation) derived from the FAMD (Fig. S11b–g).

**Fig. 3.**
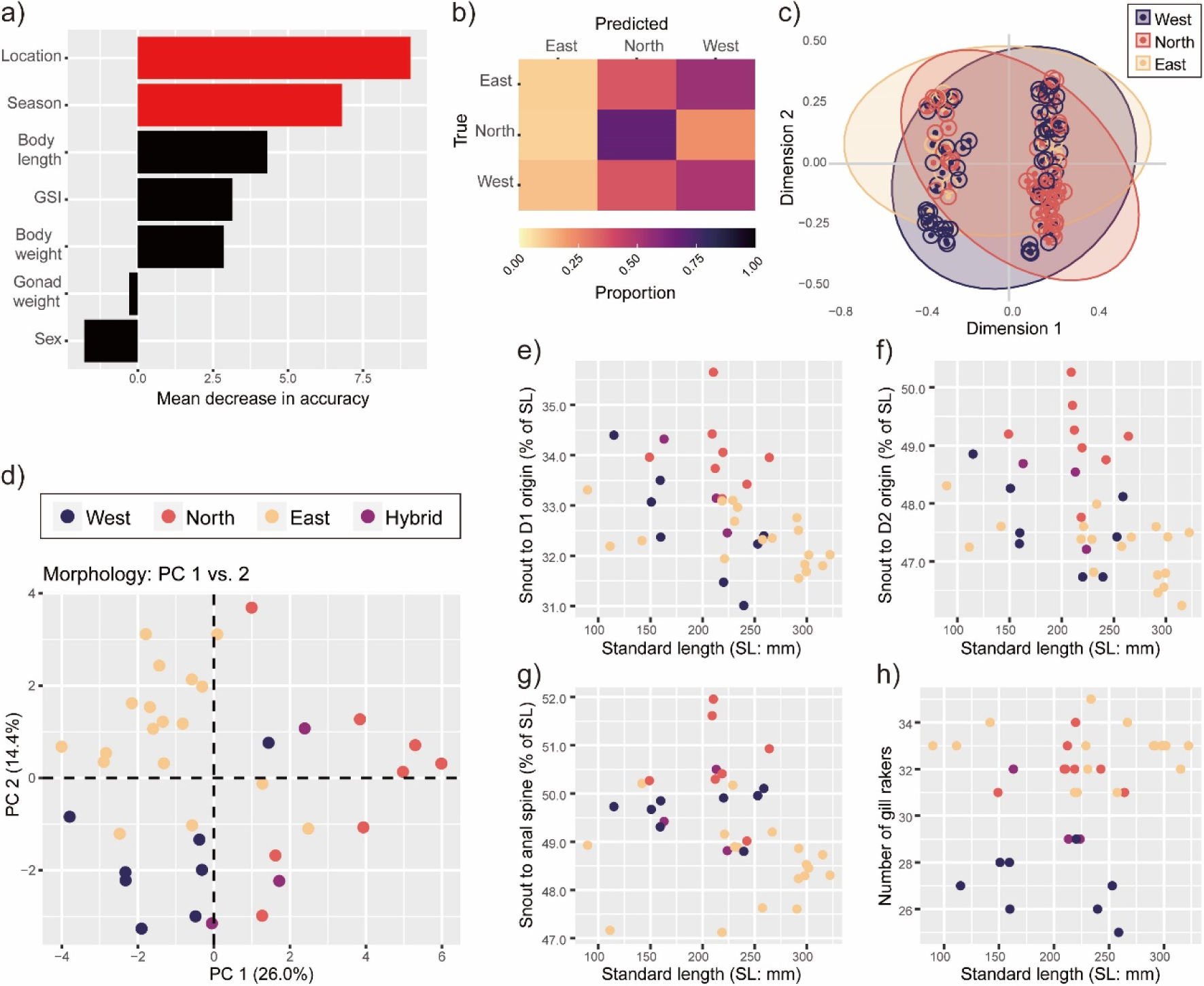
Results of the ecological (a–c) and morphological (d–h) analyses. a) Importance of ecological factors in the Random Forest model, shown by the mean decrease in accuracy. Red bars indicate significant results (permutation test, *p* < 0.05). GSI: gonad somatic index. b) Confusion matrix of the Random Forest model comparing true (genetic) individual assignments to the clusters with those predicted by the model. c) Proximity scores reduced to two multidimensional scaling dimensions, showing ecological similarity between samples as determined by the Random Forest model. Inner points represent true (genetic) assignments to the clusters, and outer circles indicate those predicted by the model. d) Principal component analysis of 24 morphological characters. The proportion of variance explained by each principal component is in parentheses. e)–h) Plots of selected morphological characters. e) The distance between the tip of the snout and the origin of the 1st dorsal fin. f) The distance between the tip of the snout and the origin of the 2nd dorsal fin g) The distance between the tip of the snout and the origin of the 1st detached anal-fin spine. h) The number of gill rakers.

### Morphology

Principal components (PCs) 1–4 explained 26.0%, 14.4%, 12.6%, and 6.7% of the total variation in overall morphology, respectively (Fig. S12a). The North cluster was separated from the East and West clusters on PC 1, suggesting its distinct morphology (Fig. 3d). This reflects the North cluster’s relatively elongated anterior body: it is distinguishable from the East and West clusters by the distances from the tip of the snout to the origins of the first-dorsal, second-dorsal, and anal fins (Fig. 3e–g), with the former two characters strongly contributing to PC 1 (Fig. S12b–d). The East and West clusters were separated from each other on PCs 2 and 3, to which gill-raker number and head width strongly contributed, respectively (Figs. 3d, S12b–e). The West cluster had fewer gill-rakers (24–29) compared to the East and North clusters (31–35 and 31–34, respectively) (Fig. 3h), implying a distinct trophic niche, as gill-raker number generally correlates with diet size and shape in fishes (e.g., Gerking 1994). Other characters did not differ considerably between the three clusters (Fig. S13). The putative hybrids occupied intermediate positions between the East and North clusters on PC 1. The discovery of distinct morphological characteristics, coupled with genetic and ecological differences, would allow us to describe the three clusters as taxonomically distinct species. See Appendix 2 for taxonomic implications.

## DISCUSSION

### Reproductive Isolation and Gene Flow

We demonstrated that the North cluster is an evolutionarily independent lineage of hybrid origin. The genome-wide differentiation (Figs. 1b, S3–S5), coupled with the presence of loci where the North cluster showed significant differentiation from both the East and West clusters (Figs. 1d, S4, S5), provides strong evidence that the North cluster is evolving separately from the others in sympatry. Eco-morphological differences further support its ecological and evolutionary independence (Figs. 3, S11–13). These results highlight that the North cluster can be recognized as a distinct species under multiple species concepts (reviewed in Coyne and Orr, 2004), such as the biological, genotypic, evolutionary, and cohesion concepts. Moreover, the genetic analyses consistently indicated the intermediate and admixed nature of the North cluster (Figs. 1c, 2a–b, S3; Tables S7–S9), and demographic model selection favored the hybrid speciation scenario over the introgression scenario (Figs. 2c, S9; Table S10). To our knowledge, this is the most compelling example of homoploid hybrid speciation in marine fishes.

**Fig. 4.**
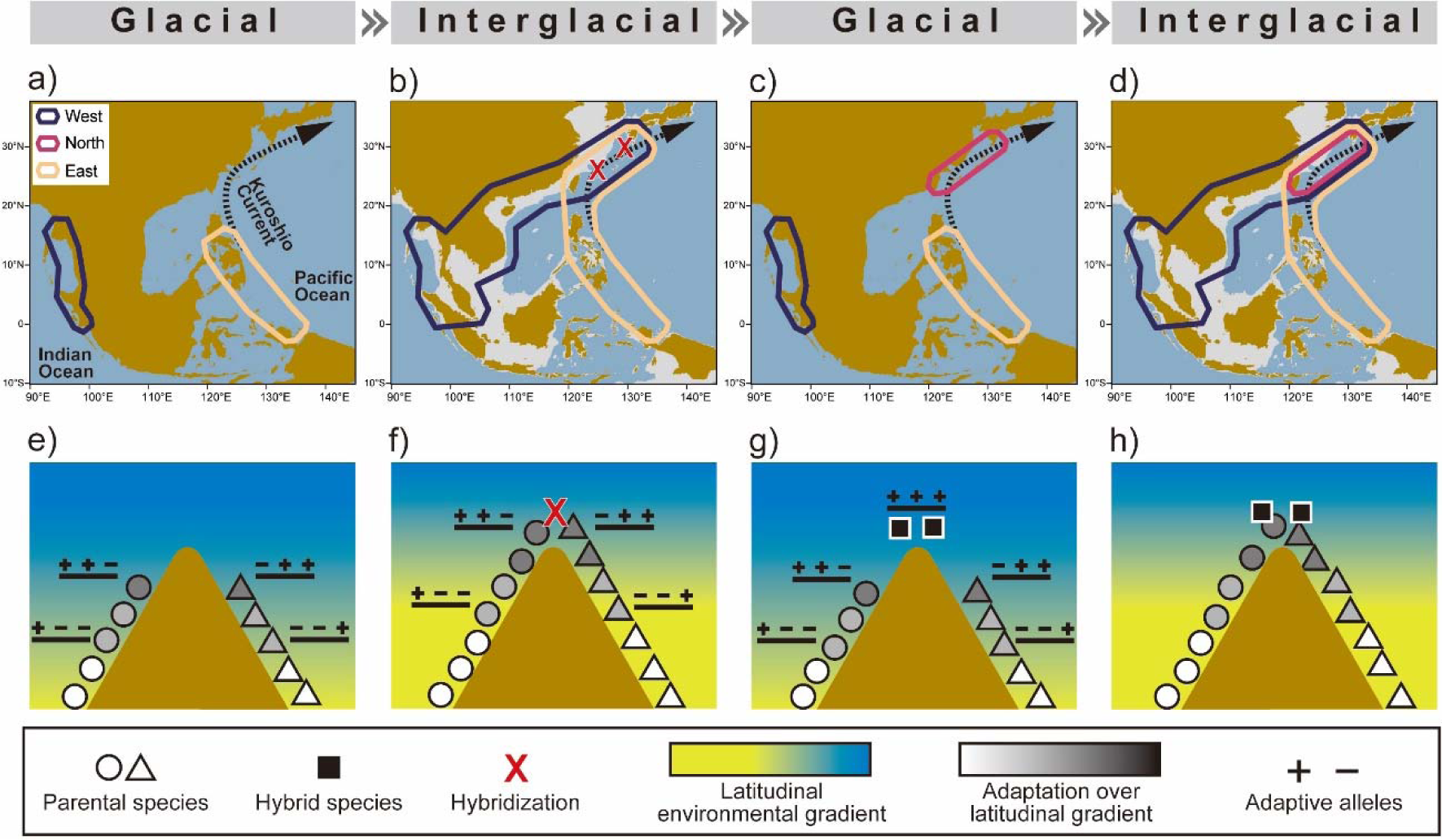
a)–d) The proposed ecogeographic scenario of HHS in the present system. Pleistocene glacial-interglacial cycles drive a) allopatric divergence of the East and West clusters, b) their northward range expansions and secondary contact, leading to the formation of the North cluster via hybridization, c) range contractions of the East and West clusters resulting in the geographical isolation of the three clusters, during which ecological and other isolating barriers develops, and d) range overlap and coexistence in sympatry. e)–h) The ecogeographic model of hybrid speciation in coastal environments driven by the Pleistocene glacial cycles. Two coastlines stretching from low to high latitudes in parallel along a landmass, joining at high-latitude extreme, are shown. Gene flow can only occur along the coastline, not across the landmass. This process can also occur where topographic features are latitudinally reversed (i.e., two coastlines join at low-latitude extreme) or the barrier to gene flow is any other uninhabitable zone, such as an open ocean as in the present system. e) During a glacial period, high-latitude coastlines become uninhabitable due to extreme conditions (e.g., low temperature), resulting in allopatric divergence of parental species. Adaptation over latitudinal gradients in temperature and other factors occurs independently, based on different loci in each species. f) Environmental change from glacial to interglacial makes the high-latitude area habitable, leading to range expansions and secondary contact of parentals, forming a hybrid population. Hybridization occurs among individuals best adapted to high latitudes since the direction of the environmental change is aligned with the latitudinal gradient. g) In the next glacial period, the hybrids have greater local fitness in high-latitude areas due to additive/epistatic effects of adaptive variants from the parentals. Additionally, heterosis in hybrids via the combination of recessive deleterious alleles, having been accumulated in the expansion front of the parentals, can also contribute the greater fitness (not depicted). The greater fitness allows hybrids to persist in high latitudes while parentals contract to low-latitude refugia, resulting in geographical isolation. During this period of allopatry, isolating barriers such as pre- and post-mating barriers based on ecological divergence may develop as by-products. h) With isolating barriers in place, the hybrid population maintains independence from parentals upon contact in subsequent interglacial periods.

The present system represents a relatively rare case of HHS where parental and hybrid species coexist in sympatry. This pattern could most readily be realized by ecological divergence between the species, which facilitates extrinsic pre- or postzygotic reproductive isolation while reducing competition (Buerkle et al. 2000). Indeed, our Random Forest analysis suggested spatio-temporal segregation of the North cluster from the others, i.e., the seasonal and locational factors strongly associated with the predictions of clusters, and the distinct number of gill-rakers in the West cluster implied its unique trophic niche (Fig. 3). We suggest that these ecological differences maintain the eco-evolutionary independence of the sympatric species. However, the ecological divergence is likely a by-product of allopatry under historical climate changes (rather than a direct outcome of hybridization), with this allopatry being the primary driver of HHS, given the estimated demographic histories and ecogeographical setting of the present system (see the next section “*Ecogeographic Scenario of Hybrid Speciation in M. cordyla*”).

It remains uncertain whether the reproductive isolation of the North cluster from the sympatric parental clusters is complete: the results of our genomic analyses were equivocal. We observed three putative F_2_s between the North and parental clusters (Table S8), indicating incomplete pre- and postzygotic isolation before the F_2_ generation. Nevertheless, postzygotic isolation due to hybrid breakdown can mainly manifest in F_2_ or later generations, effectively preventing gene flow (Orr 1995; Stelkens et al. 2015; Lollar et al. 2023). The demographic parameter estimation suggested the presence of “contemporary gene flow” (Fig. 2g); however, gene flow may be no longer ongoing, as the estimation represents an average over generations from the formation of the North cluster until the present (Fig. 2c). Another result, the uniform ancestry composition across the individuals in the North cluster (Fig. 2a–b), rather supports exclusive reproduction among the members of the same cluster, i.e., the lack of gene flow.

We observed unequal parental genomic contributions in the North cluster (Figs. 1b–c, 2a–b, S3–S5), which is typical in HHS (e.g., Elgvin et al. 2017; Barrera-Guzmán et al. 2018; Noguerales and Ortego 2022; Lopes et al. 2023; Rosser et al. 2024). Relative parental contributions in hybrid species are determined by initial contributions upon fusion, subsequent gene flow, and differential fixation of ancestry blocks via drift and selection (genomic stabilization) (Baack and Rieseberg 2007; Runemark et al. 2019). In the present system, the predominance of East cluster ancestry seems to largely reflect asymmetric gene flow following the fusion; the demographic parameter estimation suggested that the West cluster initially made a greater contribution (1−Γ_Fus_ > 0.5: Fig. 2e), and subsequent gene flow was predominantly from the East to the North cluster (Fig. 2f). Additional support for the initial closer relatedness of the West and North clusters and the importance of subsequent gene flow comes from comparing suboptimal demographic models, where the models assuming both a sister relationship between West and North (topology 2) and subsequent gene flow or pulse introgression were generally better supported than those without (Fig. S9, Table S10). The observed asymmetric gene flow may have resulted from asymmetry in the strength of reproductive isolation (Rosser et al. 2024). Alternatively, it may reflect unequal frequencies of repeated secondary contacts under the Pleistocene climatic oscillations (see the next section).

The limited number of SNPs available constrained our analytical power. This limitation led us to compromise the complexity of the demographic models, excluding events like population size changes, multiple pulses of introgression, or temporal shifts in the magnitude of gene flow, even though such events are plausible (e.g., De Jode et al. 2023; Muto and Kai 2023). Additionally, the preference for the hybrid speciation scenario over the pulse introgression was not robust, depending on the presence of both ancestral and contemporary gene flow in the model (Fig. S9). This lack of robustness might also be attributed to the limited number of SNPs. We plan a future study to address these limitations using whole genome sequencing.

### Ecogeographic Scenario of Hybrid Speciation in M. cordyla

The Pleistocene glacial cycles led to changes in sea levels and seawater temperatures, causing allopatric divergence, admixture, local adaptation, and population size changes in coastal marine organisms (Bowen et al. 2016). Based on our results and the geographic setting, we propose that multiple cycles of these processes primarily drove HHS in *M. cordyla*. Below, we outline a possible scenario in detail. Moreover, based on this scenario, we propose a model for HHS in marine coastal ecosystems, including a novel hypothesis on the initial persistence of nascent hybrid populations (see the next section “*Hybrid Speciation in Coastal Marine Environments*”).

In the Indo-West Pacific, a vast shallow continental shelf, the Sunda Shelf (Fig. 1a), strongly limited marine connectivity between the Indian and Pacific Oceans during the Pleistocene low sea-level (glacial) periods. This “Indo-Pacific barrier” drove allopatric divergence in various marine organisms, evidenced by concordant distribution patterns of sister species and phylogeographic breaks around the Sunda Shelf (Carpenter et al. 2011; Gaither and Rocha 2013). The distributions of the East and West clusters of *M. cordyla* in the tropical West Pacific align with this pattern (Fig. 1a), and their estimated divergence time fell within the Pleistocene, suggesting that they originated from the allopatric divergence across the Indo-Pacific Barrier (Fig. 4a).

Following the initial divergence, the ranges of the two clusters likely expanded and contracted in response to the glacial cycles, as demonstrated in many organisms (reviewed in Hewitt 2004; Hofreiter and Stewart 2009; Seersholm et al. 2020). Specifically, during warm interglacial periods, poleward range expansions likely occurred along mainland Asia for the West cluster and along the Kuroshio current for the East cluster (Kuriiwa et al. 2014; Motomura and Matsunuma 2022). One such poleward expansion may have resulted in their range overlap in the mid-latitude temperate area, similar to the present distribution pattern, triggering hybridization and the formation of the North cluster (Fig. 4b). In the subsequent glacial period, the North cluster survived in the temperate area with a stable population while the parentals contracted towards tropical refugia, as the Stairway Plot analysis implies (Fig. S6). This, in turn, resulted in their geographical isolation, securing the persistence of the nascent North cluster without being swamped by gene flow from the parentals (Fig. 4c). This allopatric period facilitated the development of inferred ecological difference and other unobserved isolating barriers, allowing the coexistence in sympatry during the next contact (Fig. 4d). The ancestral and contemporary gene flow inferred from the demographic modeling likely reflects relatively minor secondary contacts between the clusters.

We hypothesize that the North cluster’s survival in higher latitudes during the glacial period (i.e., Fig. 4c) was driven by clinal adaptations in the parental clusters. Prior to hybridization, parallel adaptation along latitudinal gradients in temperature and associated environmental factors likely occurred within each parental cluster, resulting in clinal patterns of adaptive genetic variants (Fig. 4e) (Jeffery et al. 2017; Teng et al. 2023). Subsequent hybridization probably took place at the high-latitude edge of the parental ranges, where these adaptive variants were most prevalent (Fig. 4f). In the North cluster, these variants recombined, enhancing fitness through additive and/or epistatic effects (Mallet 2007; Abbott et al. 2013), which enabled its survival in high latitudes during the subsequent glacial period (Fig. 4g). Conversely, deleterious alleles may have also accumulated at the expansion front of the parental clusters before hybridization, via allele surfing (Excoffier et al. 2009). However, if these deleterious alleles were recessive, they might have contributed to heterosis in hybrids, aiding their long-term persistence (MacPherson et al. 2022). Supporting our hypothesis, a recent simulation study demonstrated that admixture between two allopatric parental populations adapting to identical environments can result in hybrids that adapt more rapidly than the parentals to a novel environment (Kulmuni et al. 2024).

### Hybrid Speciation in Coastal Marine Environments

Hybrid populations can express transgressive phenotypes via combinations of parental genetic variants, exploiting novel niches and thereby becoming geographically isolated from the parental populations (Rieseberg et al. 2003; Gross and Rieseberg 2005; Mallet 2007; Abbott et al. 2013; MacPherson et al. 2022; Kulmuni et al. 2024). Although this process may underlie HHS in various organisms (Mavárez and Linares 2008; Larsen et al. 2010; Brennan et al. 2012; Kang et al. 2013; Lamichhaney et al. 2018; Maier et al. 2019; Noguerales and Ortego 2022), the ecogeographic settings enabling the process are largely unknown. The scenario described in the preceding section illustrates how this process can occur in the ocean, including the necessary ecogeographic setting. We propose this scenario as a general model of hybrid speciation in coastal marine environments driven by the Pleistocene glacial cycles (Fig. 4e–h). This model provides realistic, testable explanations for how parental adaptive genetic variants are created and how hybrids’ distinct niches lead to geographical isolation, filling the gaps in the current understanding.

Under the model, hybrid speciation can occur where coastal marine habitats in two parallel latitudinal arrays (e.g., coastlines or archipelagos) in a relevant geographical scale, separated by an uninhabitable zone (e.g., land or open ocean) but joined at latitudinal extremes (Fig. 4e). The uninhabitable zone prevents gene flow between parental species, and hybridization occurs at the latitudinal extreme under glacial maxima/minima (Fig. 4f). The following regions not only fit these topographic conditions, but also are characterized by secondary contact driven by the Pleistocene glacial cycles: temperate to tropical Indo-west Pacific (Gaither and Rocha 2013); the Japanese Archipelago (Hirase 2022); the Aleutian Archipelago (Liu et al. 2012); southern Africa (Gaither et al. 2015); and the Florida peninsula (Soltis et al. 2006). We predict that these regions harbor undiscovered cases of marine hybrid speciation.

Another key component of the model is the adaptation of the nascent hybrid population to environmental conditions at the latitudinal extreme, leveraging the preexisting adaptative genetic variants of the parents established through parallel adaptation over a common latitudinal gradient (Fig. 4f–g). The parent’s parallel adaptation may be based on similar phenotypes, but their genetic architectures are most likely *not* identical (e.g., Elmer and Meyer 2011; Jones et al. 2012), due to the lack of gene flow. Therefore, the recombinant hybrid genome can express transgressive phenotypes, conferring greater fitness than parents (Mallet 2007; Abbott et al. 2013). This greater fitness leads to local persistence under subsequent global cooling/warming and eventual allopatry from the parents, during which isolating barriers develop as by-products (Fig. 4h). These can be tested by investigating the eco-evolutionary forces underlying inter- and intraspecific genomic and phenotypic variations in parental and hybrid populations. For instance, Wang et al. (2021) employed a genome-wide association study, genomic cline analysis, and examinations of correlations between climate and genotype to investigate persistent hybrid populations of the *Setophaga* warbler complex. They demonstrated that these hybrids are differentiated from parental populations in genomic regions related to climatic adaptation, and that ancestry in these regions covaries with climatic variation.

## ACKNOWLEDGEMENTS

We thank the following persons who facilitated the sampling and data collection: H. Motomura and members of The Kagoshima University Museum (KAUM) Fish Team; H-C Ho and J-F Huang (National Museum of Marine Biology and Aquarium, Taiwan); W-J Chen and M-Y Lee (National Taiwan University); M. Nakae and K. Matsuura (National Museum of Nature and Science, Japan); T. Nakabo, Y. Kai, and M. Matsunuma (Kyoto University Museum); S. Kimura (Mie University); K. Koeda (University of the Ryukyus); M. I. Mahyam, R. H. Raja Bidin, O. Muda, W. M. Arshaad, M. Katoh, and O. Abe (Marine Fishery Resources Development and Management Department of Southeast Asian Fisheries Development Centre: SEAFDEC); Y. G. Seah (Universiti Malaysia Terengganu: UMT); A. Arshad (Universiti Putra Malaysia: UPM); S. Arnupapboon and K. Phuttharaksa (Eastern Marine Fisheries Research and Development Centre: EMDEC); N. D. The, P. V. Chien, D. V. Nhan, V. V. Chien, and P. T. Thu (Institute of Marine Environment and Resources, Vietnam: IMER); R. F. M. Traifalar, S. S. Garibay, late U. B. Alama, A. C. Gaje, and R. Cruz (University of the Philippines Visayas: UPV); Satoshi Ishikawa and members of the ‘Coastal Area Capability Enhancement in Southeast Asia’ project (Research Institute for Humanity and Nature: RIHN). We also thank Y. Kumazawa, F. Hibatullah, K. N. A. Kholil (Nagoya City University), W. Kusuma (Universitas Brawijaya), and J. R. Glass (University of Alaska Fairbanks) for their insightful discussions. The Philippines samples were collected under a Memorandum of Agreement for joint research made by and among the Department of Agriculture of the Republic of the Philippines (DA), UPV, KAUM, RIHN, and Tokai University, facilitated by S. L. Sanchez [Bureau of Fisheries and Aquatic Resources (BFAR), DA]. P. J. Alcala (DA) provided a Prior Informed Consent Certificate, and I. P. Cabacaba and S. M. S. Nolasco (BFAR, DA) provided a Fish Specimen Export Certificate (no. 2016-39812). The Malaysian samples were collected during the Japan Society for the Promotion of Science (JSPS) Asian Core Program “Establishment of Research and Education Network on Coastal Marine Science in Southeast Asia” and the JSPS Core-to-Core Program B “Asia-Africa Science Platforms” supported by UPM, UMT, and the Ministry of Higher Education of the Government of Malaysia. The Thai samples were collected with support from the EMDEC, the Training Department of SEAFDEC, and the Office of Natural Science, National Science Museum, Pathum Thani. The Vietnamese samples were collected with the support of the IMER and the Ha Long Bay Management Department and with permission to use the samples from the Biodiversity Conservation Agency, Ministry of Natural Resources and Environment, Hanoi. Computations were performed, in part, on the NIG supercomputer at ROIS National Institute of Genetics, Japan.

## FUNDING

This work was supported by the Japan Society for the Promotion of Science (JSPS) Grants-in-Aid for Scientific Research (No. JP26840131, 19K23691, 19KK0190, 23H02242, and 24K02087), JSPS Overseas Research Fellowships (No. 202160519), Sumitomo Fund (No.173246), Interdisciplinary Collaborative Research Program of Atmosphere and Ocean Research Institute, The University of Tokyo (No. JURCAOSIRG24-08), fund for integrated research activities (I.8b, 47 sub-project) from the Ministry of Agriculture and Rural Development, Vietnam, and Vietnam Academy of Science and Technology’s Grant in aid (VAST 06.04/18-19, NVCC 23.04/20-20).

## CONFLICT OF INTEREST

The authors declare no competing interests.

## AUTHORS’ CONTRIBUTIONS

NM conceived and designed the study; NM, HH, NVQ, VV, MAG, and RPB conducted sampling and sample preparation; NM conducted genetic analysis; HH and NM conducted morphological analysis; SYC conducted ecological analysis; NM wrote the manuscript with input from HH and SYC; all co-authors contributed to improving the manuscript.

## DATA AVAILABILITY

Data will be available on an online data repository upon publication of the peer-reviewed version of this manuscript.

## Notes

### Competing Interest Statement

The authors have declared no competing interest.

### Summary of Updates

Some typos were corrected and author affiliations updated.

