## Supplementary figures for "Homoploid Hybrid Speciation in a Marine Pelagic Fish *Megalaspis cordyla* (Carangidae)"

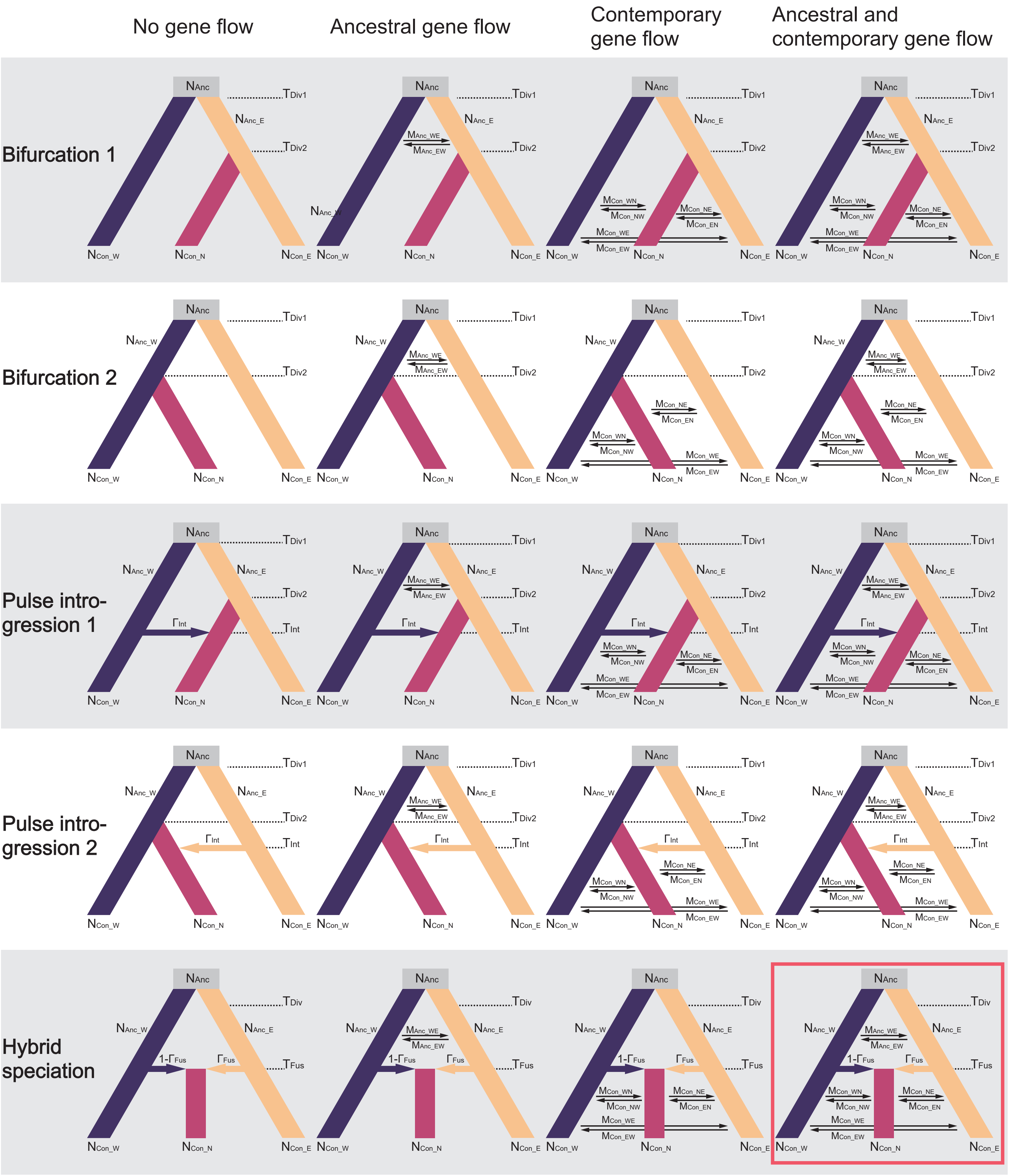


Fig. S1. Schematic diagrams of 20 demographic models tested using Fastsimcoal 2. Models include bifurcation, pulse introgression, and hybrid speciation, each tested with or without ancestral and contemporary gene flow. Two alternative topologies were assumed for the bifurcation and pulse introgression models. The best-fit model is indicated by a red rectangle. Parameters include N: ancestral ($_{Anc}$) or contemporary ($_{Con}$) effective population size of the common ancestor ($_{\_A}$) or the three genetic clusters ($_{\_E}$ / $_{\_W}$ / $_{\_N}$); T: time of divergence ($_{Div}$), fusion ($_{Fus}$), or pulse introgression ($_{Int}$) between the clusters; $\Gamma_{Fus}$: ancestry contribution from the East cluster upon fusion; 1-$\Gamma_{Fus}$: ancestry contribution from the West cluster upon fusion; $\Gamma_{Int}$: ancestry contribution from the West (topology 1) or East (topology 2) cluster upon pulse introgression; M: Ancestral ($_{Anc}$) or contemporary ($_{Con}$) gene flow between clusters, with the letters following the underscore indicating the direction of the gene flow (e.g., $M_{Con\_EW}$ represents contemporary gene flow from the East to West cluster forward in time).

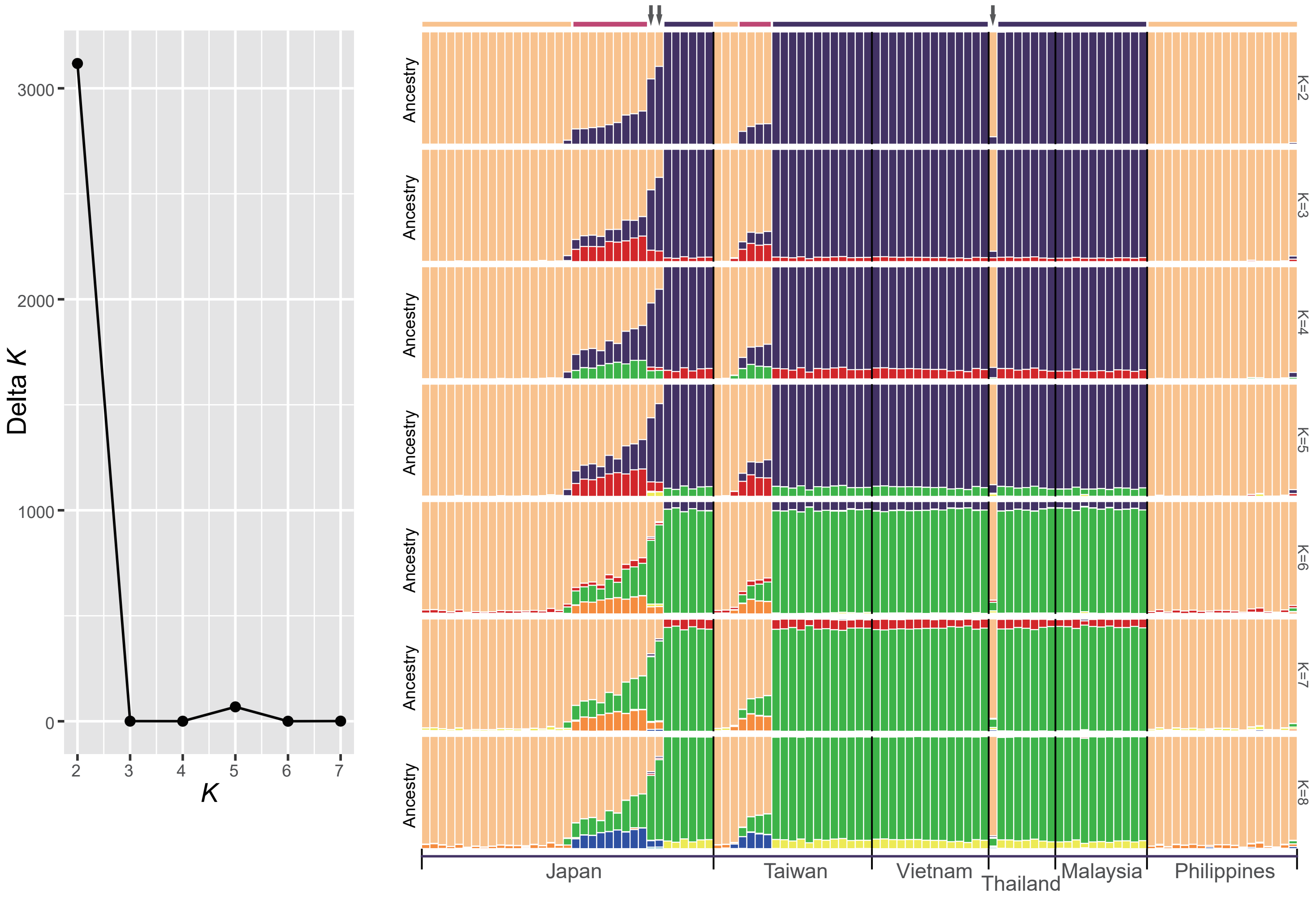


Fig. S2. Results of the STRUCTURE analysis assuming the number of clusters ($K$) = 2–7. Delta $K$ values indicate $K$ = 2 as the most likely number. Horizontal bars and arrows above the plot show individual assignments to the three clusters and putative early-generation hybrids, respectively, based on the PCA (Fig. 1b). The color code for horizontal bars matches the polygons in the PCA plot. For $K$ = 2, the North cluster and putative early-generation hybrids show admixed ancestry. For $K$ = 3 and beyond, the North cluster shows contributions from a minor cluster as well as the East and West clusters, with putative early hybrids intermediate between the North and East or West clusters.

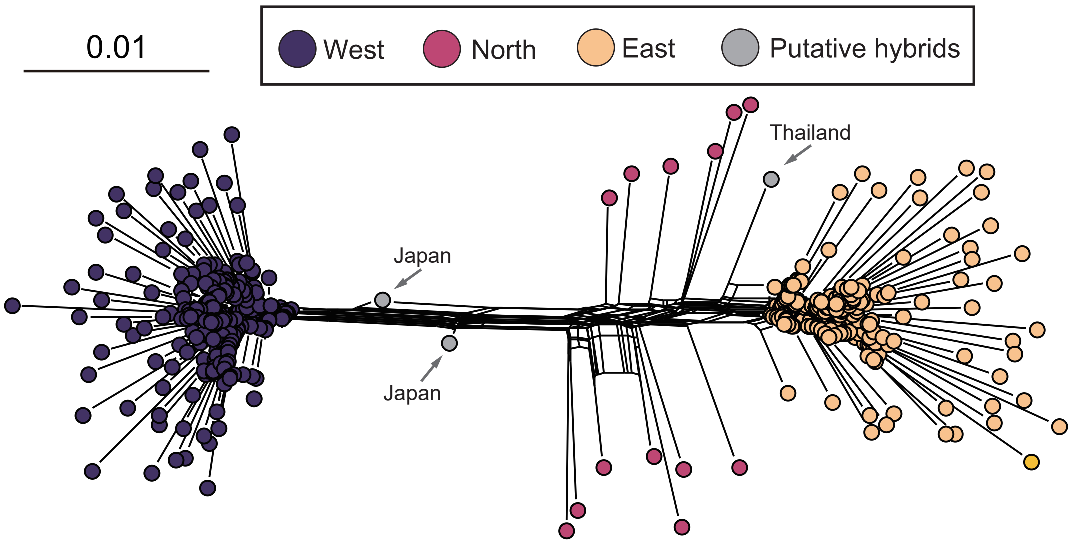


Fig. S3. A Neighbor-Net phylogenetic network inferred with SplitsTree. Individual assignments to the three clusters and putative early-generation hybrids are based on the PCA (Fig. 1b). Branch length is in units of substitutions per site.

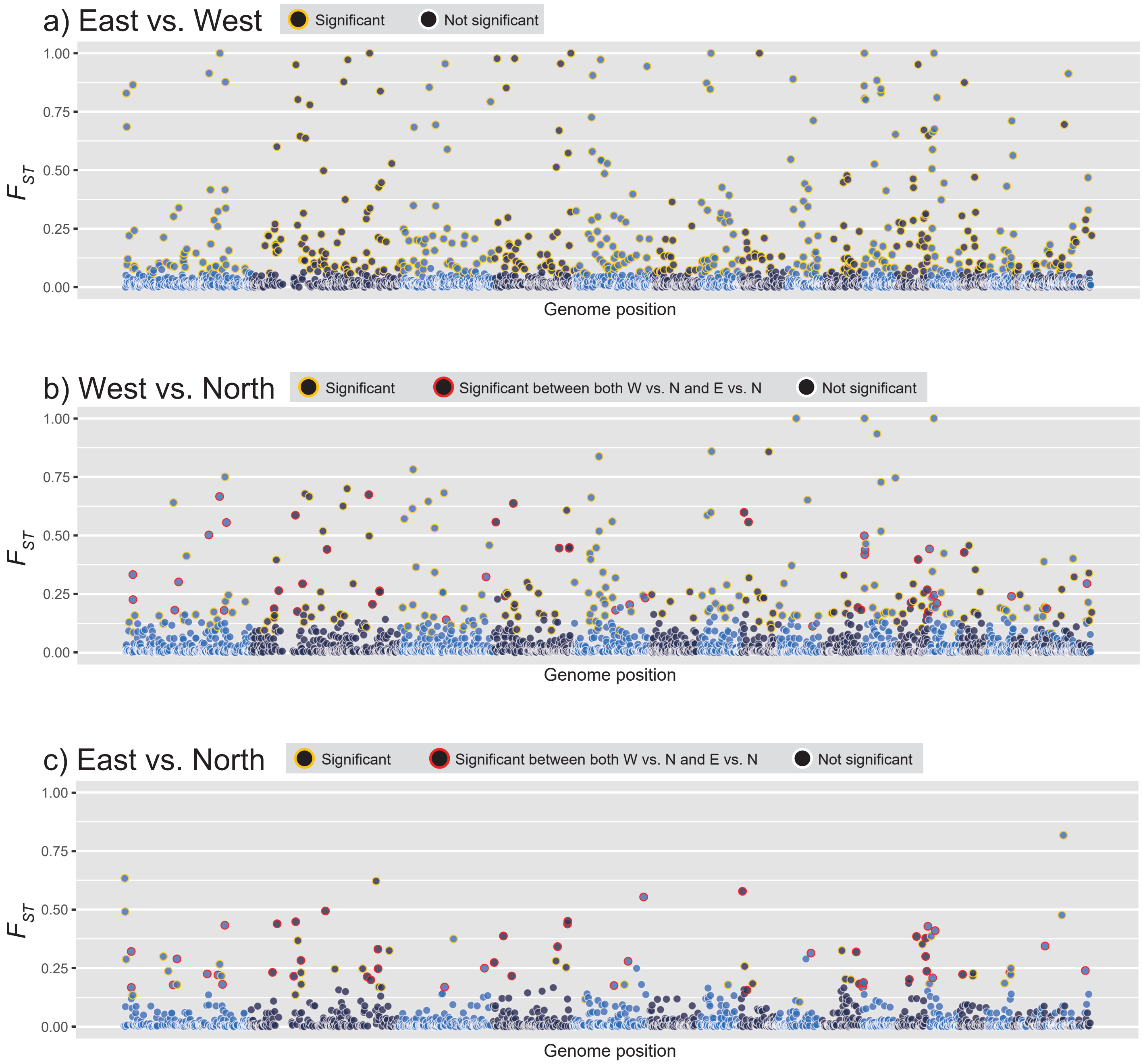


Fig. S4. Locus-specific $F_{ST}$ of 2,926 unlinked SNPs between genetic clusters, mapped to the *C. ignobilis* reference genome. SNPs on different scaffolds are shown in alternate colors. a) East-West comparison: significantly differentiated SNPs ($p < 0.01$ after false discovery correction for multiple comparisons) are indicated by a yellow margin. For the b) West-North comparison: significantly differentiated SNPs are indicated by a yellow margin; those also significant in the East-North comparison are indicated by a red margin. c) East-North comparison: significantly differentiated SNPs are indicated by a yellow margin; those also significant in the West-North comparison are indicated by a red margin.

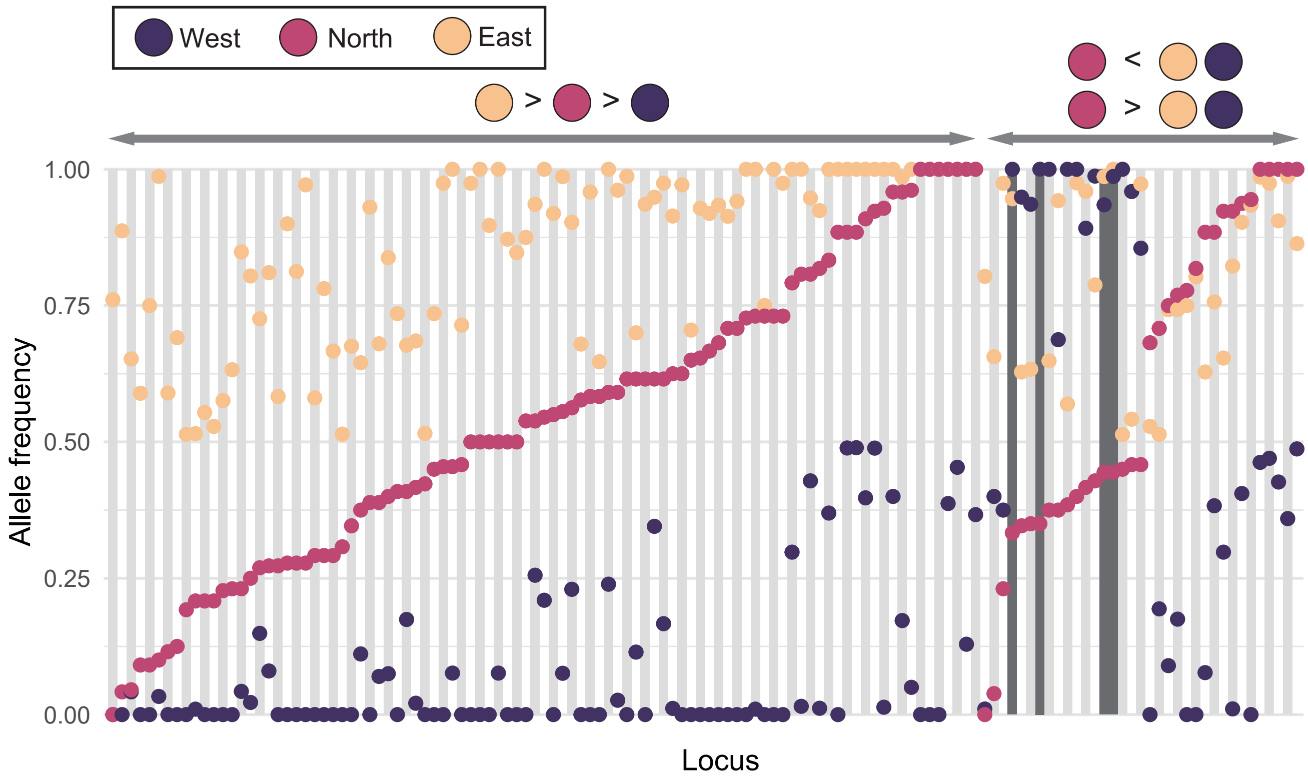


Fig. S5. Allele frequency at 130 highly differentiated sites with > 0.5 allele frequency difference between one or more pairs of clusters. The frequency of the major allele in the East cluster is shown; thus, allele frequency is > 0.5 in the East cluster by definition. The 95 sites where the North cluster's allele frequency is intermediate between the East and West clusters are grouped on the left, while the 35 sites where the North cluster's allele frequency is higher or lower than both the East and West clusters are grouped on the right. Within each of these two groups, sites are arranged in ascending order of allele frequency in the North cluster. Four sites where the North cluster's allele frequency is more than 0.5 higher or lower than both the East and West clusters are shaded dark gray.

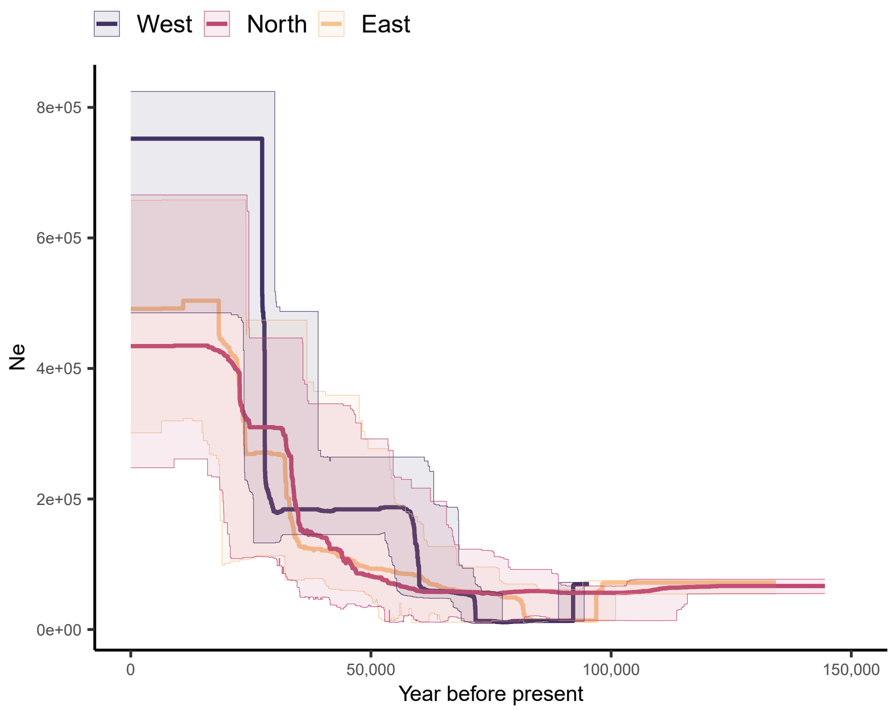


Fig. S6. Historical population size changes inferred by Stairway Plot 2. The thick line represents the median; the thin lines represent the 95% confidence intervals.

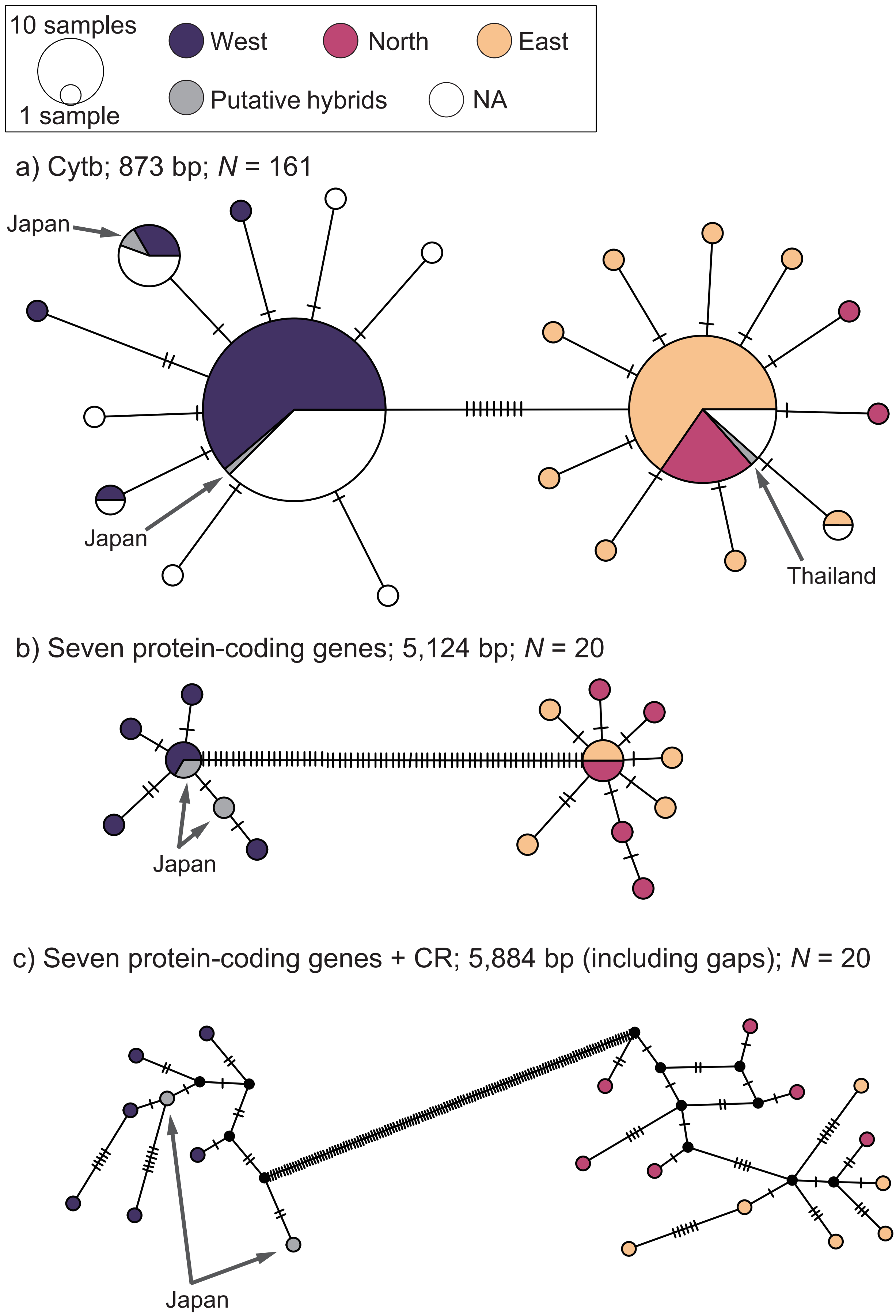


Fig. S7. Statistically parsimonious networks of haplotypes for a) CYTB sequences, b) concatenated sequences of 7 protein-coding mitochondrial genes (CYTB, COI, COII, ND1, ND2, ND4, and ND5), and c) concatenated sequences of the 7 protein-coding genes plus the control region. Circle size is proportional to haplotype frequency. Black dots and hatch marks represent unobserved hypothetical haplotypes and one mutational step, respectively. Individual assignments to the three clusters and putative early-generation hybrids are based on the PCA (Fig. 1b). "NA" indicates the specimens without SNPs data.

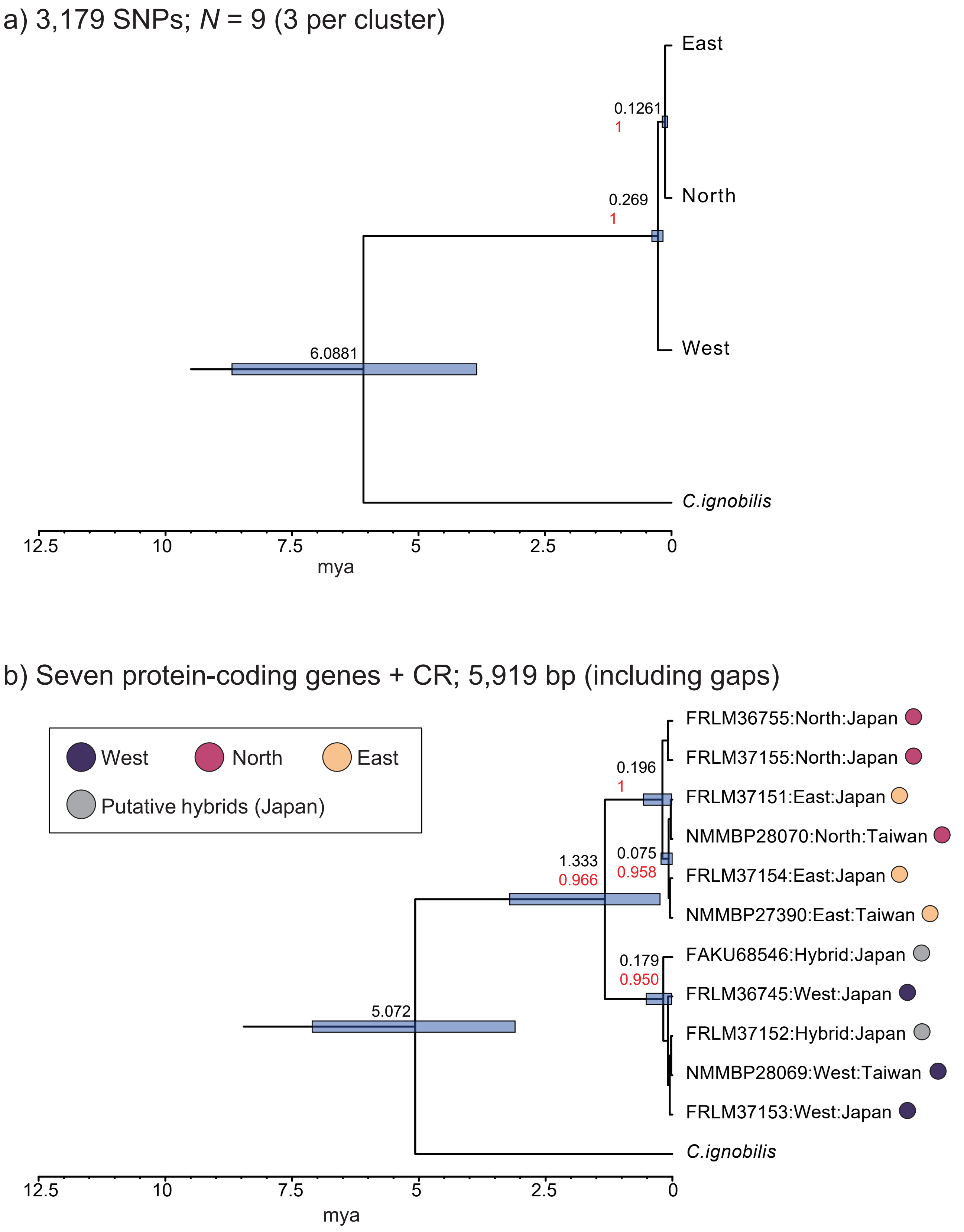


Fig. S8. a) A multispecies coalescent species tree inferred with SNAPP based on 3,179 unliked SNPs and b) mtDNA phylogeny inferred with Beast 2 based on the concatenated sequences of the 7 protein-coding genes plus control region. OTUs are genetic clusters in a) and individuals in b). Numbers beside nodes represent mean age (upper row: black) and posterior probability (lower row: red), with bars representing 95% highest posterior density intervals of node age. Only nodes with posterior probability > 0.95 are shown.

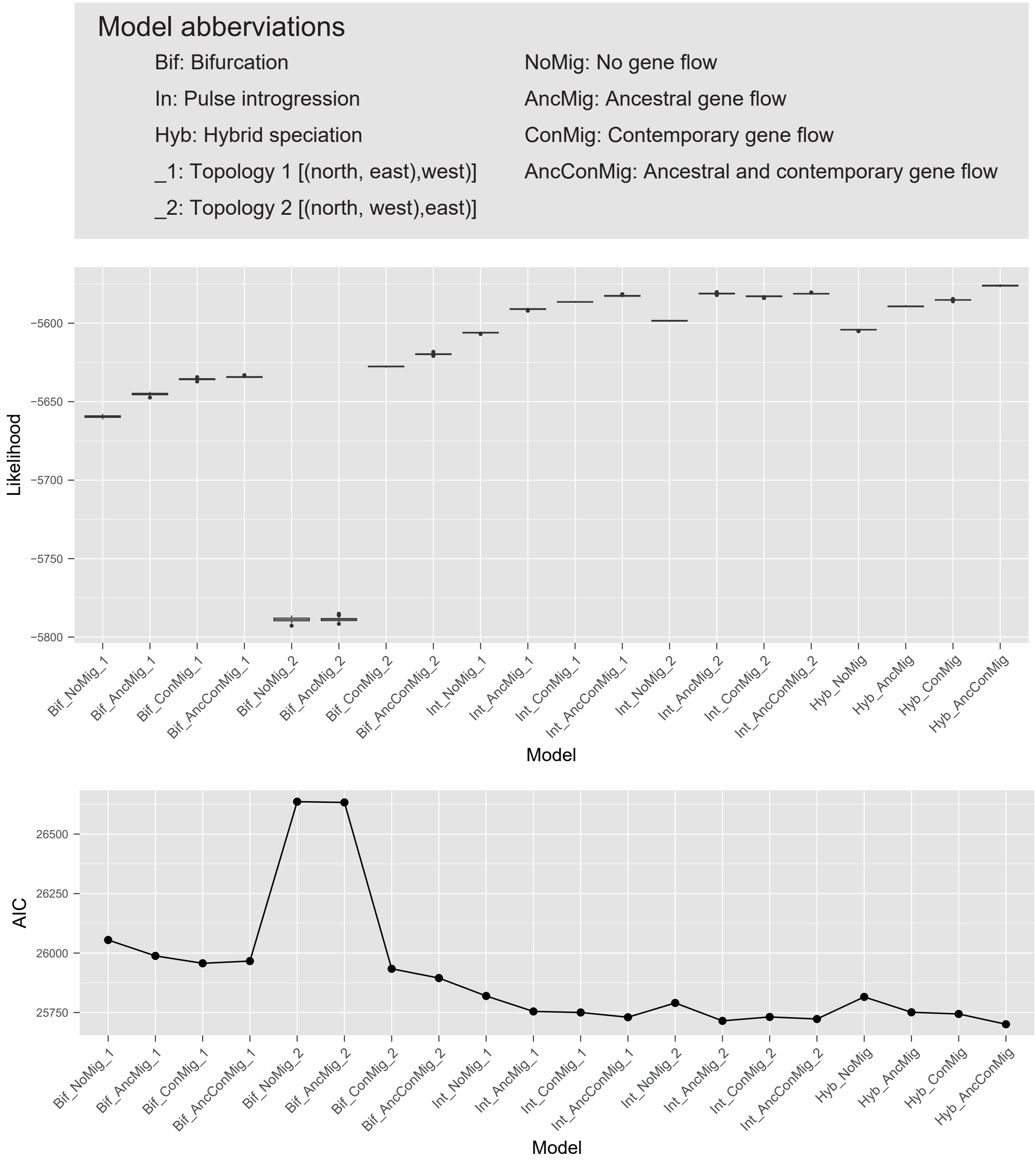


Fig. S9. Likelihood distributions and Akaike’ s information criterion (AIC) values for 20 alternative demographic models.

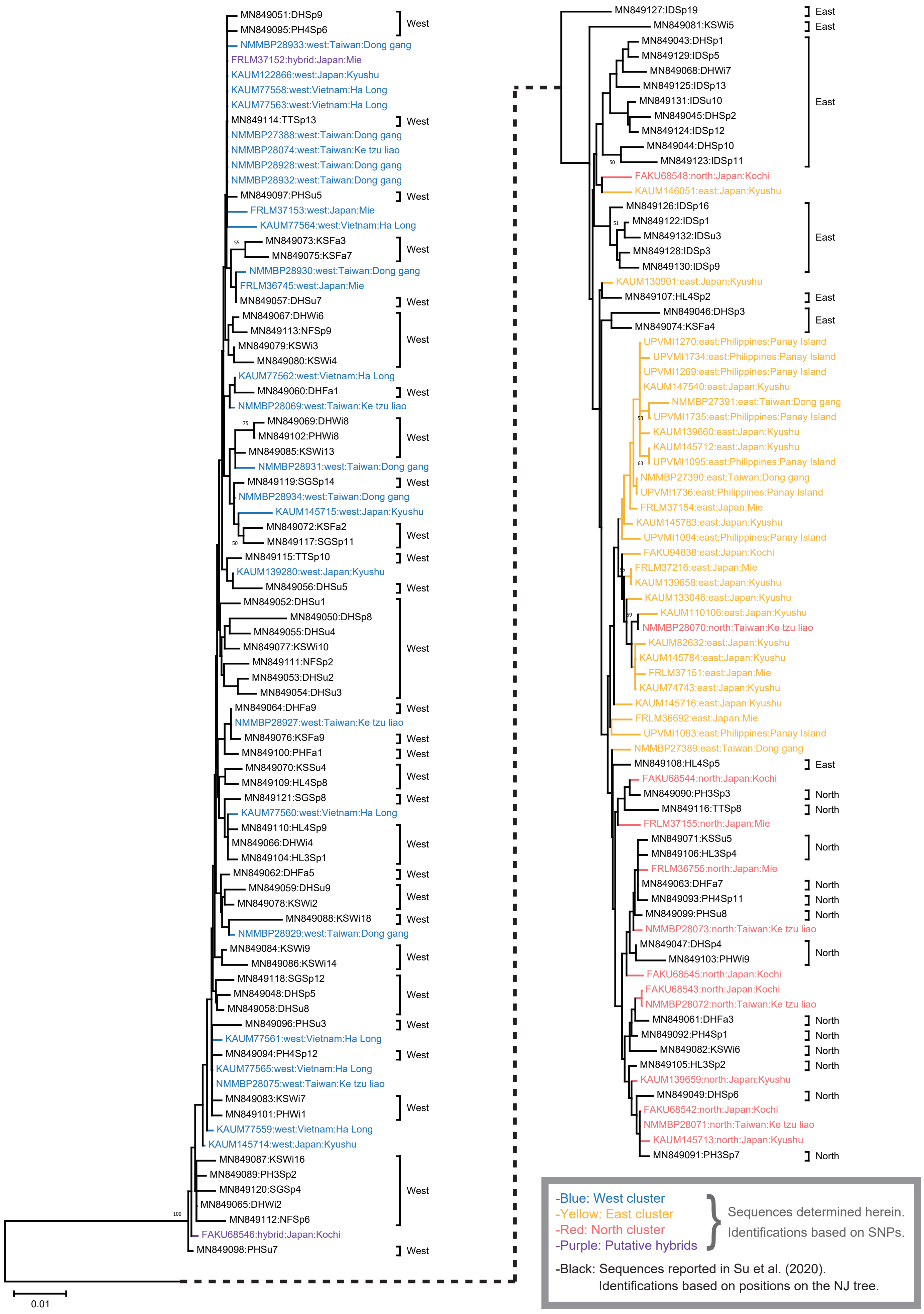


Fig. S10. A Neighbor-Joining tree based on a 592 bp alignment of the mtCR sequences for identifications of the haplotypes reported by Su et al. (2020) to the three genetic clusters discovered herein. Identification is shown by text next to each haplotype based on its position on the NJ tree relative to sequences from the present specimens. The present specimens have been identified to the three genetic clusters (indicated by color) by the genome-wide SNPs genotyping.

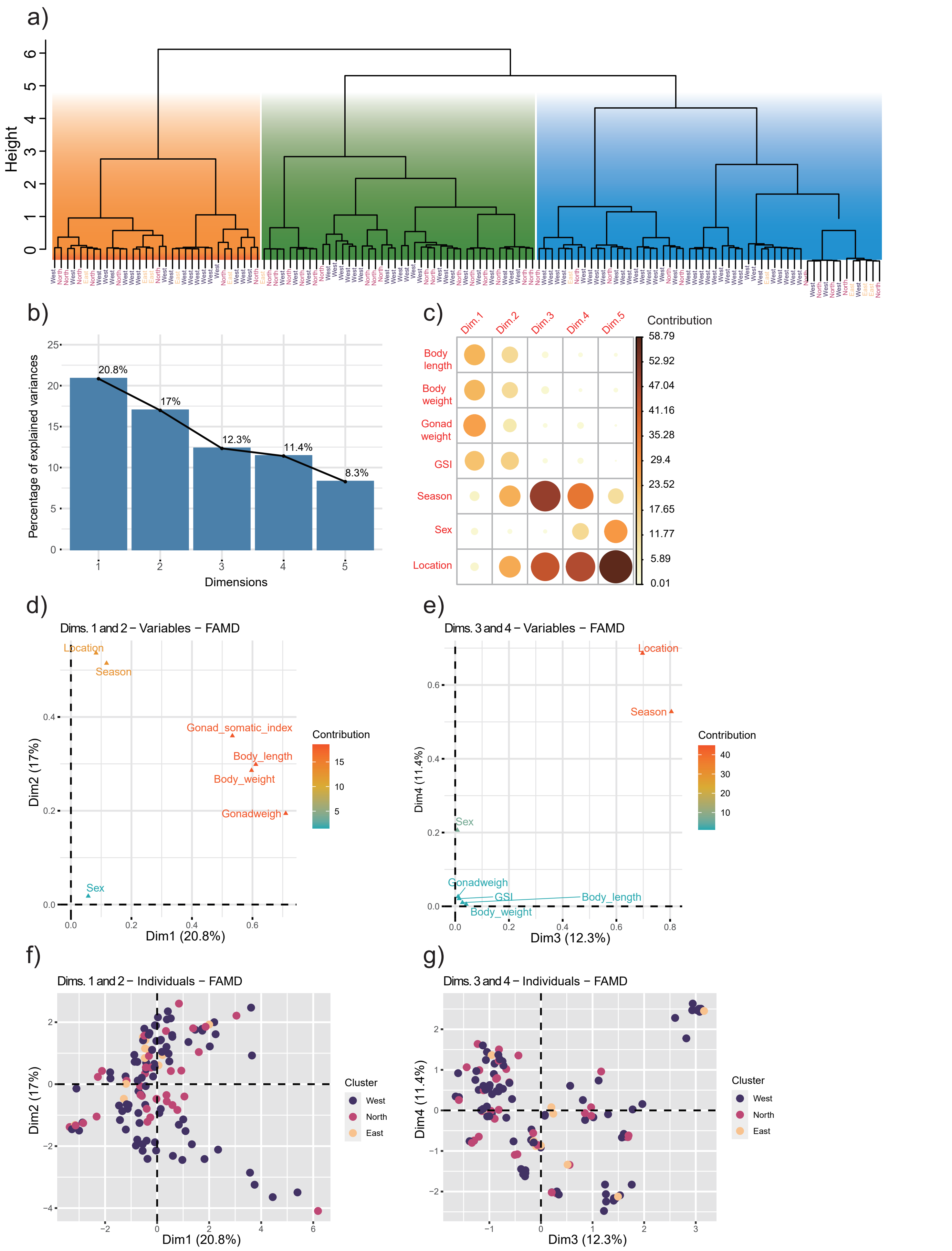


Fig. S11. Results of the a) hierarchical cluster analysis and b–g) factor analysis of mixed data (FAMD) of the ecological data of *M. cordyla* in Taiwan reported by Su et al (2020). a) The dendrogram obtained through the hierarchical cluster analysis. Terminal nodes represent samples having been analyzed by Su et al (2020) and herein re-identified to the three genetic clusters according to the mtCR sequences (Fig. S10; Table S4). Rectangles indicate clusters in the dendrogram recognized under an arbitrary cutoff of the number of clusters = 3. b) Percentages of variance explained by the first 5 dimensions of FAMD. c) Contributions of the ecological factors to the 5 dimensions. d), e) Coordinates of the ecological factors on the dimensions, with colors indicating their contributions. f), g) Coordinates of individual samples on the dimensions.

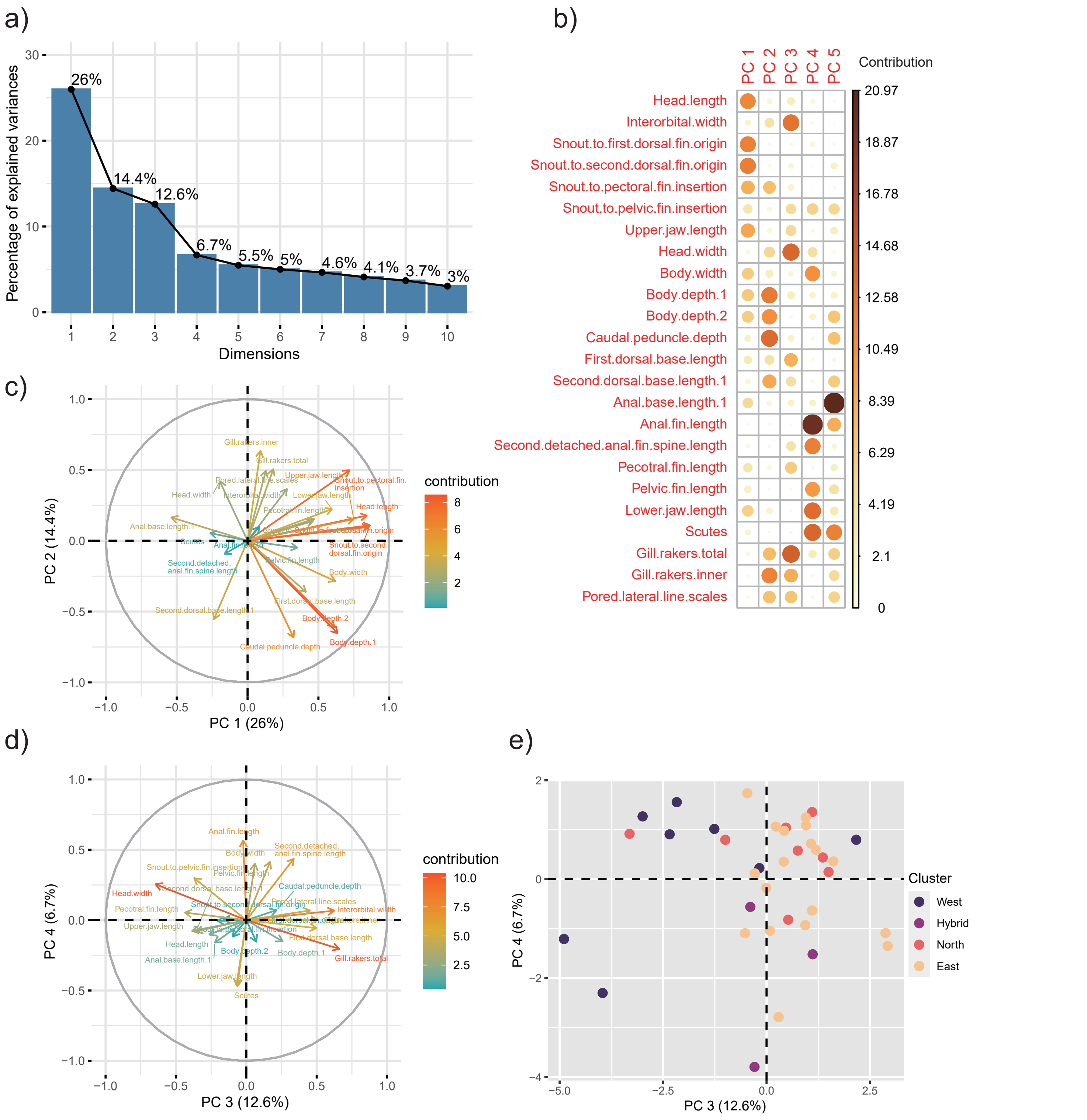


Fig. S12. Results of the principal component analysis (PCA) of 24 morphological characters. a) Percentages of variance explained by the first 10 PCs. b) Contributions of the morphological characters to the first 5 PCs. c), d) Coordinates of the morphological characters on the PCs, with colors indicating their contributions. e) Coordinates of individual samples on Pcs 3 and 4.

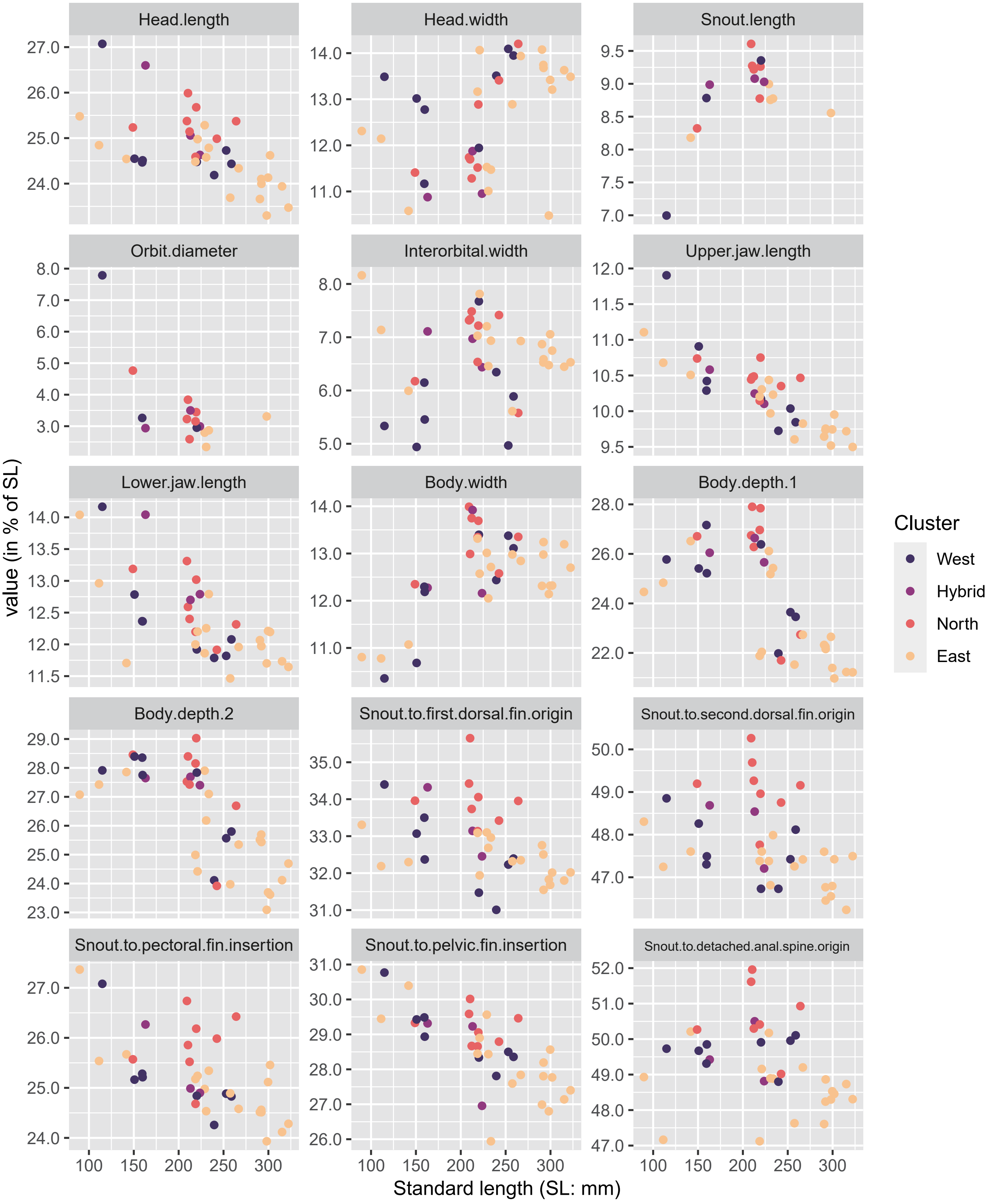


Fig. S13. Plots for all morphological characters examined.

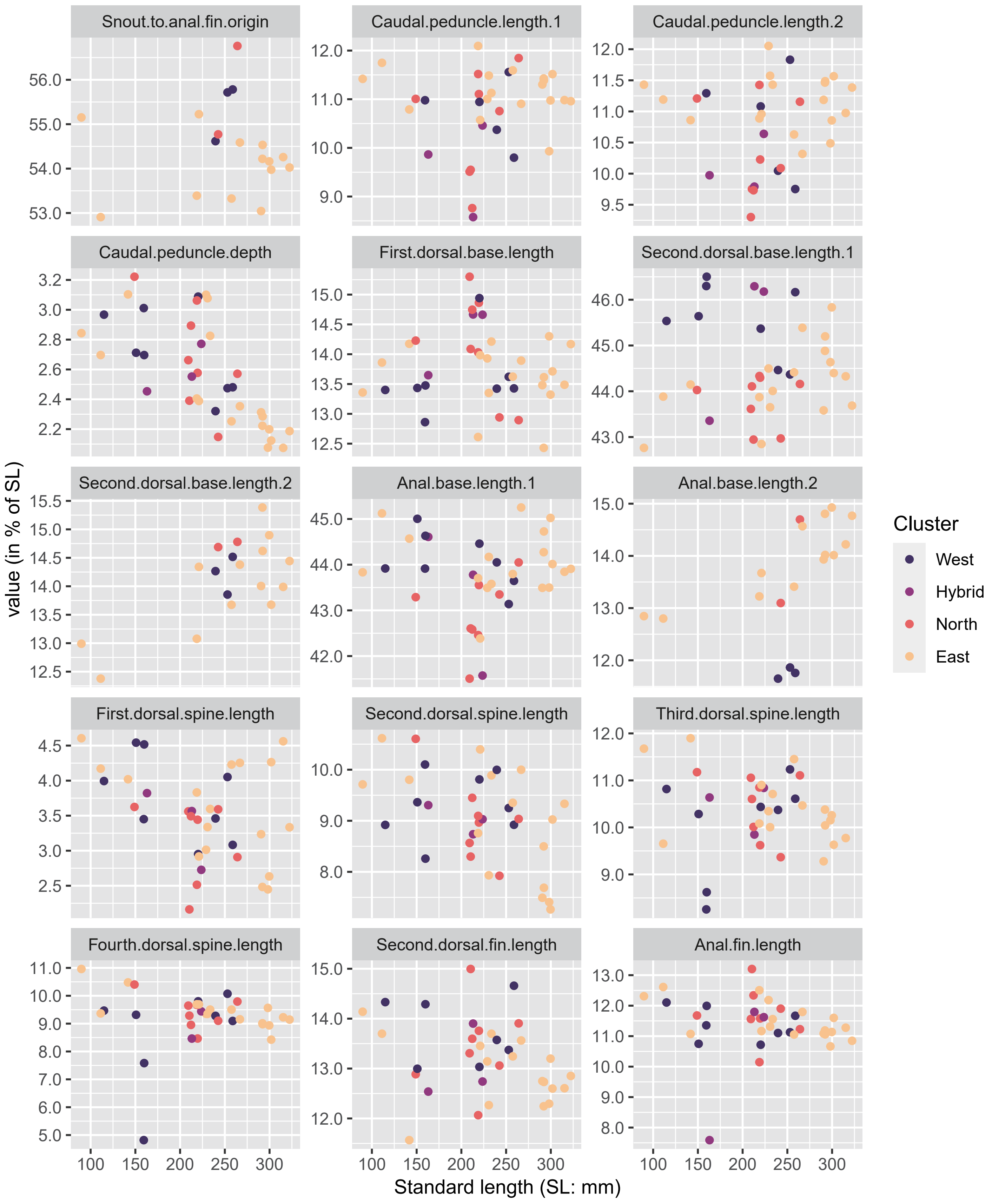


Fig. S13 (continued)

Fig. S

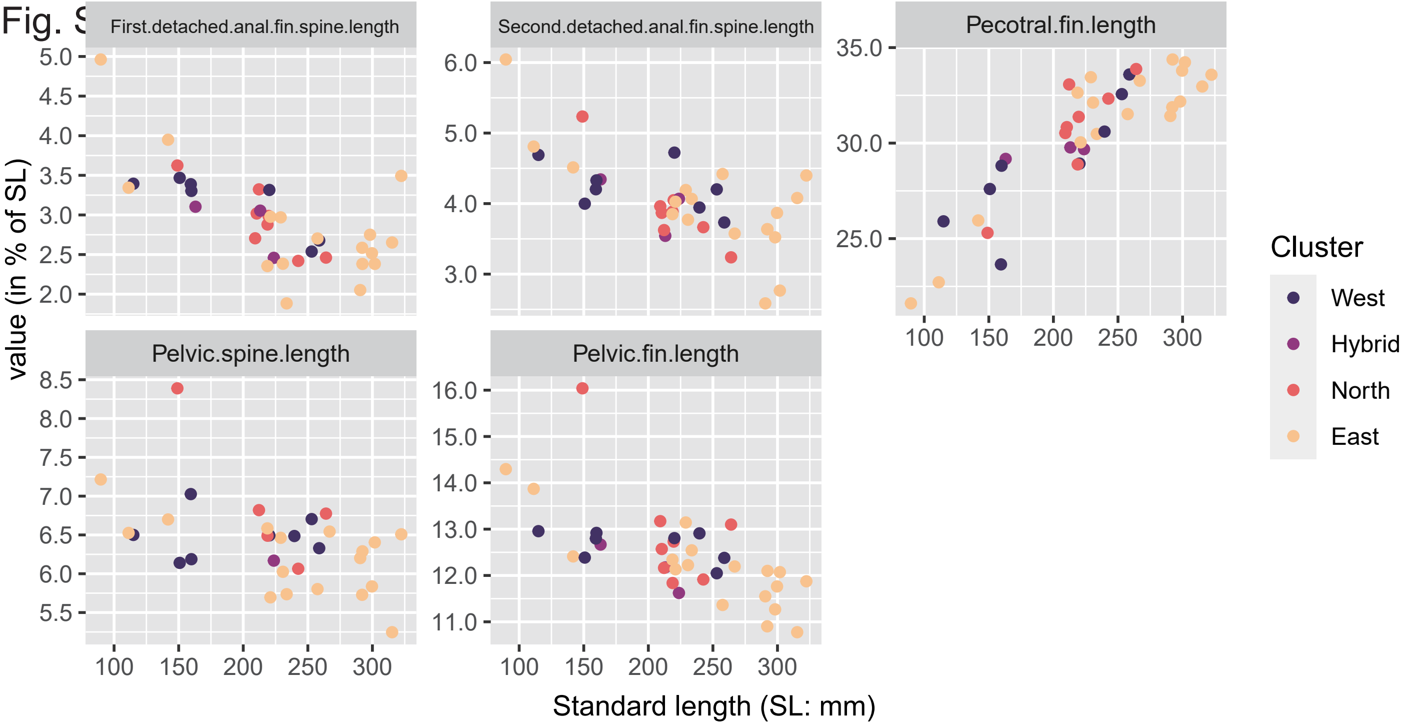


Fig. S13 (continued)

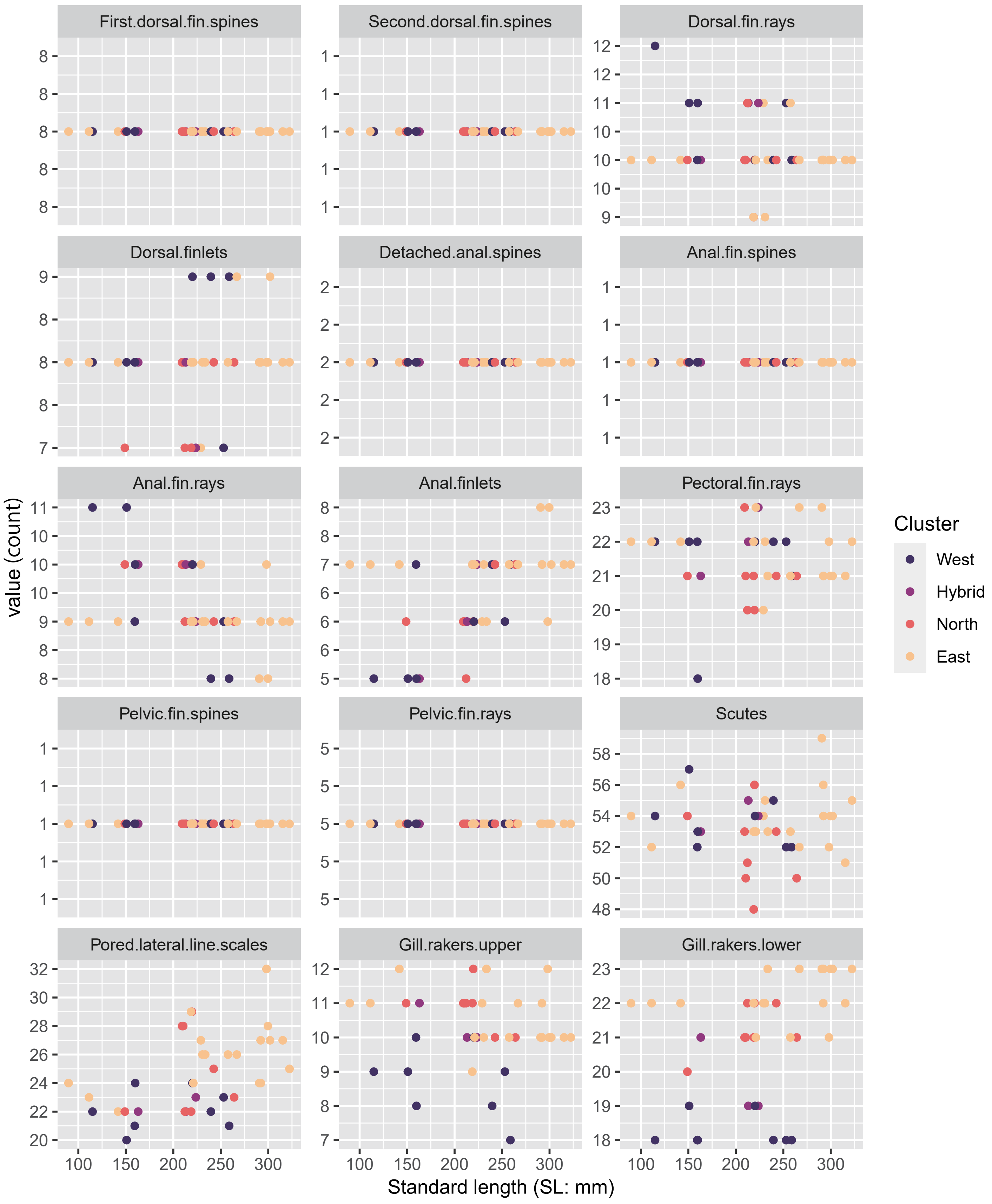


Fig. S13 (continued)

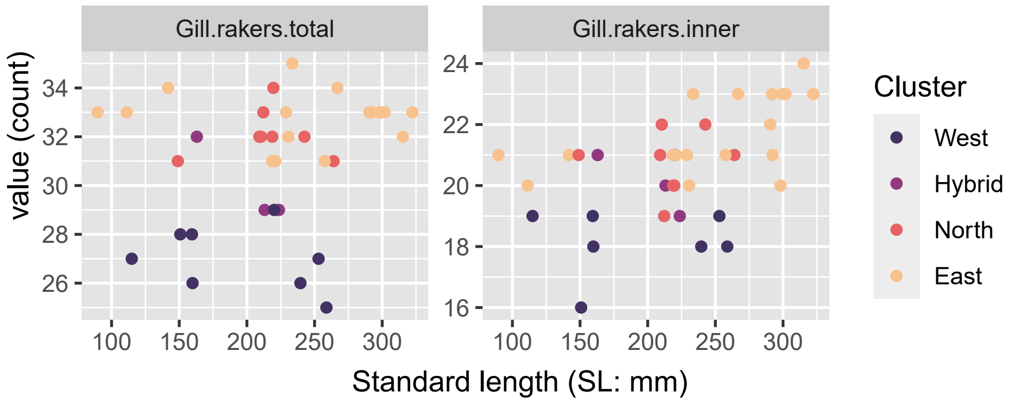


Fig. S13 (continued)
