## Appendix for "Homoploid Hybrid Speciation in a Marine Pelagic Fish *Megalaspis cordyla* (Carangidae)"

**Appendix 1: Full description of methods**

*Mitochondrial DNA Genotyping*

We obtained the partial sequences of the mitochondrial cytochrome b gene (CYTB: 873 bp) from all specimens. Additionally, we obtained the partial sequences of the following 7 loci from subset of the specimens: cytochrome c oxidase subunits 1 (COI: 612 bp) and 2 (COII: 657 bp), NADH dehydrogenase subunits 1 (ND1: 828 bp), 2 (ND2: 531 bp), 4 (ND4: 828 bp), and 5 (ND5: 801 bp) and the control region (CR: 760 bp) (Table S1). We amplified the COI sequence using a universal primer cocktail comprising four (two each of forward and reverse) primers: VF2_t1, FishF2_t1, Fish R2_t1, and FR1d_t1 (‘COI-3’) (Ivanova et al. 2007). We amplified the CYTB sequence using the CTB-F and CTB-R primers (Cárdenas et al. 2005). The remaining loci were amplified using newly developed primers (Table S2). We performed PCR in a total volume of 8.0 µL containing 4.0 µL of KAPA2G Robust HS ReadyMix with dye (KAPA Biosystems), 0.125 µM of each of the four primers (for COI) or 0.25 µM of each of the forward and reverse primers (for the other genes), and 1.0 µl (approximately 10–100 ng) of template DNA. Thermal profiles for COI and CYTB followed the aforementioned original publications. Thermal profiles for the other loci are shown in Table S2. PCR products were purified using ExoSAP-IT (Thermo Fisher Scientific) according to the manufacturer’s protocol. DNA sequencing was performed using the Big-Dye Terminator Cycle Sequencing Kit v3.1 (Thermo Fisher Scientific) on an ABI 3500 Genetic Analyzer (Thermo Fisher Scientific). We edited and aligned the obtained sequences using BioEdit v7.0.5.3 (Hall 1999) and CLUSTAL X (Larkin et al. 2007), respectively.

*Genome-Wide SNPs Genotyping*

We genotyped 115 specimens using MIG-seq (Suyama and Matsuki 2015; Suyama et al. 2022), a reduced representation sequencing method targeting SNPs in the inter-simple sequence repeat region (ISSR). We constructed three MIG-seq libraries consisting of PCR products of 24, 64, and 67 specimens. The first library consisted of 24 specimens from Japan and Taiwan. These 24 specimens were sequenced twice in the first and third libraries to supplement the insufficient amount of data from the first sequencing. The second library contained 16 specimens that were not used in the present study. The libraries were sequenced using an Illumina Novaseq 6000 in 150 bp paired-end mode at Novogen Co.Ltd., China, yielding 8.5, 58.6, and 80.9 M reads, respectively. We performed the first PCR in a total volume of 10.0 µL containing 5.0 µL of KAPA2G Fast Multiplex Mix (KAPA Biosystems), 0.2 µM of each of the 12 multiplex primers (Suyama and Matsuki 2015; Suyama et al. 2022), and 1.0 µl of template DNA, with the following thermal profile: 95°C for 3 min, followed by 25 cycles at 95°C for 15 sec, 38°C for 1 min, 72°C for 1 min, and a final extension at 72°C for 10 min. We performed the second PCR in a total volume of 15.0 µL containing 7.5 µL of KAPA HiFi HS ReadyMix (KAPA Biosystems), 0.2 µM of each primer designed to incorporate illumina sequencing adapters and a unique dual index for each specimen, and 3.0 µl of the first PCR product that was diluted 50-folds. The thermal profile for the second PCR was: 95°C for 3 min, followed by 20 cycles at 98°C for 20 sec, 65°C for 15 s, 72°C for 30 s, and a final extension at 72°C for 5 min. We pooled the second PCR products in equivalent volumes, and the resulting library was purified with MinElute PCR Purification Kit (QIAGEN), size-selected for a target range of 400–800 bp using SPRIselect (BECKMAN COULTER), and then purified again using GeneRead Size Selection Kit (QIAGEN). Subsequently, the quality of the libraries was checked using a Quantus Fluorometer (Promega) and an Agilent 4200 TapeStation system (Agilent).

We processed both reads 1 and 2 of the raw MIG-seq reads by the single-end mode of fastp v0.20.1 (Chen et al. 2018) to remove the first 17 bases containing simple sequence repeat, adapter sequences, and low-quality reads, and to trim the reads to 100 bp, with the following parameters: -f 17 -b 100 -q 20 -l 100. We merged the quality-filtered reads 1 and 2 of each specimen into a single FASTQ file and mapped them to the reference genome assembly of *Caranx ignobilis* (GenBank JAFHLA000000000.1) (Pickett et al. 2022) using BWA-mem v0.7.17 (Li 2013) with default settings. Unmapped reads and reads with a mapping quality of < 20 were removed using Samtools v1.12 (Danecek et al. 2021). The resulting BAM files were used to prepare 5 datasets for downstream analyses (see the next paragraph). We also downloaded the whole-genome shotgun sequencing reads of *C. ignobilis* (different individual from the one used for the reference genome assembly) and *C. melampygus* from the NCBI Short Read Archive (accession nos.SRR1002875 and SRR1002876) (Santos 2014) for use as outgroups. We first processed these reads using the paired-end mode of fastp to remove adapter sequences, low-quality reads, and short reads (< 25 bp) with the following parameters: -q 20 -l 25. We mapped the processed reads to the reference genome of *C. ignobilis* using BWA-mem with default settings. Unmapped reads, reads with a mapping quality of < 20, and duplicate reads were removed using Samtools. The resulting BAM files were used for the analyses requiring outgroups (datasets 2–5: see below).

The following 5 datasets were prepared to meet requirements or preferable conditions for a variety of downstream analyses: 1) 2,926 unlinked biallelic SNPs in all available (105) specimens of *M. cordyla* without an outgroup, used for the population genetic analyses using various tools and the ancestry estimation using HIest v2.0 (Fitzpatrick 2012); 2) 6,789 unlinked biallelic SNPs in 24 specimens (8 specimens per genetic cluster) with *C. ignobilis* as an outgroup, for the admixture detection using HyDe v1.0.0 (Blischak et al. 2018) and ADMIXTOOLS v7.0.2 (Patterson et al. 2012); 3) 3,179 unlinked biallelic SNPs in 9 specimens (3 per genetic cluster) with *C. ignobilis* as an outgroup, for the divergence time estimation using SNAPP v1.5.0 (Bryant et al. 2012); 4) 6,341 biallelic SNPs in the same 24 specimens, polarized using the sequences of *C. ignobilis* and *C. melampygus* as outgroups, for the demographic model selection using fastsimcoal v2.7.0.9 (Excoffier et al. 2013, 2021); and 5) 22,326 biallelic SNPs (both linked and unlinked) and 920,184 monomorphic sites in the same 24 specimens as above, polarized using the sequences of *C. ignobilis* and *C. melampygus* as outgroups, for the demographic parameter estimation using fastsimcoal 2 and the historical population size estimation using Stairway Plot v2.1.2 (Liu and Fu 2020). Note that the monomorphic sites and linked SNPs included in the 5th dataset are necessary for direct parameter estimations using fastsimcoal 2, whereas only unlinked SNPs are used for the model selection (Excoffier et al. 2013). For the 1st dataset, we called SNPs and created a vcf file using *gstacks* and *populations* implemented in Stacks v2.5.4 (Catchen et al. 2013). We then retained only genotypes/sites that fulfilled the following criteria using VCFtools v0.1.16 (Danecek et al. 2011) and R package ‘SNPfiltR’ v1.0.0 (DeRaad 2022) on R v4.2.2 (R Core Team 2022): genotypes with read depth of > 5, allele balance of > 0.2 and < 0.8, and genotype quality of > 20; sites with two alleles (biallelic sites), overall depth across specimens < 100, genotyping rate across specimens > 0.75, and minor allele count of > 1. We further filtered the resulting sites by their positions on the reference genome so that they were separated apart at least 5 kb from each other. We then removed 10 specimens with a genotyping rate of < 0.5. For the 2nd dataset, we called genotypes for the 24 specimens (8 specimens per cluster with the highest number of mapped reads) together with *C. ignobilis* as an outgroup using *mpileup* implemented in bcftools v1.8 (Danecek et al. 2021) with default settings. We filtered the resulting genotypes/sites in the same way as the 1st dataset, except that the sites with minor allele count of = 1 were retained for the 2nd dataset. We then removed the sites that were only variable between *C. ignobilis* and *M. cordyla* (i.e., monomorphic in *M. cordyla*). For the 3rd dataset, we further reduced the number of specimens to 3 per cluster. We called genotypes and filtered the data by the same procedure as described for the 2nd dataset, except that sites that were variable only between *C. ignobilis* and *M. cordyla* were retained for the 3rd dataset. This enabled the estimation of the divergence times between the genetic clusters of *M. cordyla* relative to that between *C. ignobilis* and *M. cordyla*. We also thinned the sites so that they were separated apart at least 100 kb (instead of 5 kb for the 1st and 2nd datasets) from each other to overcome the high computational demand of SNAPP. For the 4th dataset, we called genotypes and filtered the resulting data in the same way for the 2nd dataset, and retained only sites that passed two additional filtering criteria: 1) fixed for the reference allele in both outgroups (*C. ignobilis* and *C. melampygus*) and 2) biallelic within *M. cordyla*. Finally, the 5th dataset was prepared in the same way as the 4th dataset except that the sites were not filtered by their positions on the reference genome, and that monomorphic sites were retained.

We used the *easySFS.py* script (https://github.com/isaacovercast/easySFS) to calculate the unfolded SFS from the 4th and 5th datasets, assuming that the reference alleles fixed in both outgroups represent the ancestral state. We selected the projection with the maximum number of segregating sites, which was 12 for each cluster for both datasets. The resulting one-dimensional SFS based on the 5th dataset was used for the historical population size estimation by Stairway Plot 2, whereas multi-dimensional SFS was used for the demographic inference by fastsimcoal 2. For the value to be input to the monomorphic cell in the multi-dimensional SFS based on the 5th dataset (required for the parameter estimation), we calculated the ratio of monomorphic and polymorphic sites in the entire dataset and multiplied the ratio by the total number of SNPs in the multidimensional SFS following (Ruffley et al. 2022). For the preparation of input files for the other analyses, we used PGDSpider v2.1.1.2 (Lischer and Excoffier 2012) to convert the vcf files to appropriate formats.

*Population Genetic Analyses*

We constructed minimum-spanning haplotype networks of mtDNA sequences using PopART v1.7 (Leigh and Bryant 2015).

The following population genetic analyses were conducted using 1st dataset of the genome-wide SNPs. We performed a principal component analysis (PCA) using the ‘adegenet’ package v2.1.4 in R (Jombart 2008). We used STRUCTURE v2.3.4 (Pritchard et al. 2000) to examine the presence of admixed individuals. We performed 10 replicate runs for a predefined number of genetic clusters (*K*) = 1–8, each consisting of 100,000 Markov Chain Monte Carlo (MCMC) generations after 100,000 generations of burn-in. These runs were parallelized using StrAuto v1.0 (Chhatre and Emerson 2017). Statistical support for the different *K* was evaluated by calculating the Δ*K* value (Evanno et al. 2005) using Structure Harvester v0.6.94 (Earl and vonHoldt 2012). The results were summarized and visualized with ‘pophelper’ v2.3.1 in R (Francis 2017). We used SplitsTree v5.3.0 (Huson and Bryant 2006) to build an unrooted phylogenetic network based on the Neighbor-Net algorithm.

The population genetic analyses described above showed that one of the genetic clusters was genetically intermediate to the other two clusters (Fig. 1b, c). The intermediate cluster is hereafter called the “North” cluster, whereas the others are called “East” and “West” clusters, according to their geographical distributions (Fig. 1a). Moreover, putative early-generation hybrids (F1, F2, or backcross) between these clusters were detected. Excluding these putative early-generation hybrids, we calculated locus-specific pairwise *F_ST_* between clusters using *populations* implemented in Stacks and pairwise *F_ST_* across all loci and associated *p*-values using Arlequin v3.5.2.2 (Excoffier and Lischer 2010).

We estimated historical population size changes in each cluster using Stairway Plot v2.1.2 (Liu and Fu 2020) based on dataset 5. We assumed a mutation rate of 1E^−8^ per site per generation (Tine et al. 2014) and a generation time of 1.2 years derived from FishBase (Froese and Pauly 2023). The generation time of *M. cordyla* (1.2 years) was approximated as the age class with the maximum egg production (t_opt_), which was calculated using the parameters of the von Bertalanffy growth function as t_opt_ = t_0_ − ln(1 − L_opt_ / L_inf_) / K, where t_0_, L_opt_, L_inf_, and K represent the hypothetical age the fish would have had at zero length, the length class of giving highest biomass, the asymptotic length, and the rate at which the asymptotic length is approached, respectively. For *M. cordyla* these parameters are estimated as: t_0_ = −0.20; L_opt_ = 31.6; L_inf_ = 49.6; and K = 0.71. The other settings of Stairway Plot 2 were set to default values.

*Divergence Time Estimation*

We estimated the divergence times of the three genetic clusters based on the genome-wide SNPs dataset 3 and mtDNA sequence data. The former estimation was done under the multispecies coalescent model using the SNAPP v1.5.0 package (Bryant et al. 2012) of BEAST v2.7.1 (Bouckaert et al. 2014), following (Stange et al. 2018). Time calibration was based on the estimated node age of 6.349 (95% CI: 4.434-8.698) Ma for the most recent common ancestor (MRCA) of *C. ignobilis* and *M. cordyla* (Glass et al. 2023). This age calibration was approximated by a lognormal distribution with a mean of 6.349 Ma and a standard deviation of 0.205 and an offset of -0.01. We performed two independent runs of 50,000,000 MCMC generations, sampling every 10,000 steps. We used Tracer v1.7.1 (Rambaut et al. 2018) to check the convergence of the chain and confirm that effective sample sizes (ESS) for all parameters were > 200. We discarded the first 10% of the trees as burn-in, and the posterior tree distributions from the two replicates were combined using LogCombiner v2.7.1 (Bouckaert et al. 2014). We estimated a maximum clade credibility tree using TreeAnnotator v2.7.1 (Bouckaert et al. 2014) and visualized the tree using FigTree v1.4.4 (http://tree.bio.ed.ac.uk/software/figtree/). The estimation based on the concatenated sequences of the 8 mtDNA loci was done using BEAST v2.7.1. We partitioned the sequences by locus and estimated the substitution model for each partition using the bModelTest package v1.3.3 (Bouckaert and Drummond 2017) of BEAST. The bModelTest package averages over different substitution models implemented in BEAST, rather than relying on a single model. For all partitions, we assumed the optimized relaxed clock model and the Yule model for the clock model and the tree prior, respectively. We used the mitogenome sequence of *C. ignobilis* (accession No.: NC_022932) as an outgroup. Time calibration, MCMC simulations, convergence check, tree estimation, and visualization were the same as described above.

*Ancestry Estimation and Admixture Detection*

We estimated the ancestries of the individuals in the North cluster and the putative early-generation hybrids using the ‘HIest’ package v2.0 (Fitzpatrick 2012) in R. This program estimates the ancestries of target individuals relative to predefined parents via log-likelihood maximization, using genotypes of the targets and allele frequencies of the parents. We estimated the ancestries of the individuals in the North cluster and the putative early-generation hybrids, assuming the East and West clusters to be potential parents and using 108 unlinked SNPs with allele frequencies differing by > 0.5 between them. We also estimated the ancestries of putative hybrids, assuming the three clusters as potential parents, using 130 SNPs with allele frequencies differing by >0.5 between one or more pairs of clusters. In the first analysis, we estimated the ancestry index (*S*) and the inter-class heterozygosity (*H*) using the *HIest* function, employing the SANN method for searching the likelihood surface, 500 MCMC generations, and the option surf = TRUE. *S* represents the proportion of an individual’s alleles derived from one of the two predefined parental populations (in the present case, the East cluster), equivalent to the hybrid index. *H* indicates the proportion of an individual’s loci that have one allele from each parental population. Expected *H* values vary between classes of early-generation hybrids (F1: 1.0; F2 and backcross: 0.5), becoming progressively lower if succeeding reproduction occurs exclusively among hybrids and alleles are lost by drift. We also estimated the likelihoods for each of the six possible genotype classes resulting from 2 generations of inter- or intraspecific crosses (parental, F1, F2, and backcrosses) using *HIclass* function, comparing the likelihood of the best-fit of the six genotypes with that of the maximum likelihood estimation of *S* and *H* using the *HItest* function. For the second analysis, we estimated the ancestry indices *S*_East_, *S*_West_, and *S*_North_ of the putative early-generation hybrids using the *threeway* function. These indices are equivalent to *S*, representing the proportion of an individual’s alleles derived from the three predefined parental populations. We also calculated the likelihoods for each of the 15 genotype classes possible in the first two generations of hybridization between pairs of parental populations using the *thirdclass* function, comparing the maximum likelihood with that obtained from the *threeway* analysis using the *HItest3* function.

*Morphology*

We examined 47 linear measurements, 24 countable characters, and 3 binary characters in 37 specimens from Japan, Taiwan, Vietnam, and Thailand (Tables S1, S3). This analysis aimed to determine if the genetic clusters 1) differ in ecologically relevant morphological characters, suggesting ecological divergence, and 2) can be recognized as separate species in a taxonomic sense. Linear measurements were made to the nearest 0.01 mm using a digital vernier caliper. Methods for measuring and counting generally followed (Marr and Schaefer 1949), (Gibbs and Collette 1967), and (Muto et al. 2016) except for the following―Lower-jaw length followed Hubbs and Lagler (1947); Snout to detached anal-spine origin and snout to anal-fin origin are the distances from the tip of the snout to the origin of the 1st detached anal spine and the 1st attached anal spine, respectively; Caudal peduncle lengths 1 and 2 are the distances from the posterior end of the hypural to the insertions of the last dorsal and anal finlets, respectively; Second dorsal base lengths 1 and 2 are the lengths of the 2nd dorsal-fin base excluding and including the dorsal finlets, respectively; Anal base lengths 1 and 2 are the anal-fin base lengths excluding and including both the detached anal spines and anal finlets, respectively; Scutes were counted on the straight portion of the lateral line; Pored lateral line scales were counted excluding the scutes; Gill rakers inner are the number of gill rakers on the inner side of the 1st gill arch. Linear measurements are expressed as % of the Standard length (SL). To illustrate the variation in overall morphology, we performed a PCA of 24 characters with no missing data using the R package `FactoMineR*`* v2.11 (Lê et al. 2008). All linear measurements were corrected for allometry using the R package ‘GroupStruct’ v0.1.0 (Chan and Grismer 2022) before performing the PCA.

*Ecology*

To investigate potential ecological divergence between the genetic clusters, we re-analyzed the ecological data of *M. cordyla* in Taiwan reported by Su et al. (2020). The data include sampling season and location, individual body length and weight, gonad weight, gonad somatic index (GSI: [gonad weight/body weight] × 100), and sex (Table S4), along with the mtCR sequences of the specimens. Su et al. (2020) compared the ecological factors between two major lineages of mtCR sequences. In the present study, we show that one of these two major lineages corresponded to the West cluster, while the other corresponded to the East and North clusters (see Results). Additionally, we demonstrate that the East and North clusters can be distinguished by a minor mtCR divergence within the major lineage. These observations allowed us to reclassify the specimens examined by Su et al. (2020) into the three clusters based solely on the mtCR sequences reported by the authors, albeit with some uncertainty. First, we tested the association between the ecological factors and genetic clusters under the Random Forest model using the R package ‘ranger’ v0.16.0 (Wright and Ziegler 2017). We assessed the importance of each ecological factor using the mean decrease in accuracy and estimated the significance of this importance metric with a permutation test using the R package ‘rfPermute’ v2.1.81 (Archer 2021). Second, we performed a hierarchical cluster analysis to create a dendrogram of the specimens based on the ecological data. We calculated the pairwise Gower’s distance between each specimen and constructed Ward’s hierarchical agglomerative clustering using the R package ‘stats’ v 4.2.2 (R Core Team 2022). Third, we performed a Factor Analysis of Mixed Data (FAMD) using the R package ‘FactomineR’.

*References*

Archer E. 2021. rfPermute: Estimate Permutation p-Values for Random Forest Importance Metrics. https://CRAN.R-project.org/package=rfPermute.

Blischak P.D., Chifman J., Wolfe A.D., Kubatko L.S. 2018. HyDe: A Python package for genome-scale hybridization detection. Syst. Biol. 67:821–829.

Bouckaert R., Heled J., Kühnert D., Vaughan T., Wu C.H., Xie D., Suchard M.A., Rambaut A., Drummond A.J. 2014. BEAST 2: A software platform for Bayesian evolutionary analysis. PLOS Comp. Biol. 10:1–6.

Bouckaert R.R., Drummond A.J. 2017. bModelTest: Bayesian phylogenetic site model averaging and model comparison. BMC Evol. Biol. 17:1–11.

Bryant D., Bouckaert R., Felsenstein J., Rosenberg N.A., RoyChoudhury A. 2012. Inferring species trees directly from biallelic genetic markers: bypassing gene trees in a full coalescent analysis. Mol. Biol. Evol. 29:1917–1932.

Cárdenas L., Hernández C.E., Poulin E., Magoulas A., Kornfield I., Ojeda F.P. 2005. Origin, diversification, and historical biogeography of the genus *Trachurus* (Perciformes: Carangidae). Mol. Phylogenet. Evol. 35:496–507.

Catchen J., Hohenlohe P.A., Bassham S., Amores A., Cresko W.A. 2013. Stacks: an analysis tool set for population genomics. Mol. Ecol. 22:3124–3140.

Chan K.O., Grismer L.L. 2022. GroupStruct: An R Package for Allometric Size Correction. Zootaxa. 5124:471–482.

Chen S., Zhou Y., Chen Y., Gu J. 2018. fastp: an ultra-fast all-in-one FASTQ preprocessor. Bioinformatics. 34:i884–i890.

Chhatre V.E., Emerson K.J. 2017. StrAuto: automation and parallelization of STRUCTURE analysis. BMC Bioinformatics. 18:192.

Danecek P., Auton A., Abecasis G., Albers C.A., Banks E., DePristo M.A., Handsaker R.E., Lunter G., Marth G.T., Sherry S.T., McVean G., Durbin R., 1000 Genomes Project Analysis Group. 2011. The variant call format and VCFtools. Bioinformatics. 27:2156–2158.

Danecek P., Bonfield J.K., Liddle J., Marshall J., Ohan V., Pollard M.O., Whitwham A., Keane T., McCarthy S.A., Davies R.M., Li H. 2021. Twelve years of SAMtools and BCFtools. GigaScience. 10:giab008.

DeRaad D.A. 2022. snpfiltr: An R package for interactive and reproducible SNP filtering. Mol. Ecol. Resour. 22:2443–2453.

Earl D.A., vonHoldt B.M. 2012. STRUCTURE HARVESTER: a website and program for visualizing STRUCTURE output and implementing the Evanno method. Conserv. Genet. Resour. 4:359–361.

Evanno G., Regnaut S., Goudet J. 2005. Detecting the number of clusters of individuals using the software STRUCTURE: a simulation study. Mol. Ecol. 14:2611–2620.

Excoffier L., Dupanloup I., Huerta-Sánchez E., Sousa V.C., Foll M. 2013. Robust demographic inference from genomic and SNP data. PLOS Genet. 9:e1003905.

Excoffier L., Lischer H.E.L. 2010. Arlequin suite ver 3.5: a new series of programs to perform population genetics analyses under Linux and Windows. Mol. Ecol. Resour. 10:564–567.

Excoffier L., Marchi N., Marques D.A., Matthey-Doret R., Gouy A., Sousa V.C. 2021. fastsimcoal2: demographic inference under complex evolutionary scenarios. Bioinformatics. 37:4882–4885.

Fitzpatrick B.M. 2012. Estimating ancestry and heterozygosity of hybrids using molecular markers. BMC Evol. Biol. 12:131.

Francis R.M. 2017. pophelper: an R package and web app to analyse and visualize population structure. Mol. Ecol. Resour. 17:27–32.

Froese R., Pauly D. 2023. Fishbase. Available from https://www.fishbase.se/popdyn/KeyfactsSummary_1.php?ID=384&GenusName=Megalaspis&SpeciesName=cordyla&vStockCode=398&fc=314.

Gibbs R.H., Collette B.B. 1967. Comparative anatomy and systematics of the tunas, genus *Thunnus*. Fish. Bull. 66:65–130.

Glass J.R., Harrington R.C., Cowman P.F., Faircloth B.C., Near T.J. 2023. Widespread sympatry in a species-rich clade of marine fishes (Carangoidei). Proc. R. Soc. B. 290:20230657.

Hall T.A. 1999. BioEdit : a user-friendly biological sequence alignment editor and analysis program for Windows 95/98/NT. Nucleic Acids. Symp. Ser. 41:95–98.

Hubbs C. L., Lagler K. F. 1947. Fishes of the Great Lakes Region. Bull. Cranbrook Inst. Sci. 26:1–186.

Huson D.H., Bryant D. 2006. Application of phylogenetic networks in evolutionary studies. Mol. Biol. Evol. 23:254–267.

Ivanova N.V., Zemlak T.S., Hanner R.H., Hebert P.D.N. 2007. Universal primer cocktails for fish DNA barcoding. Mol. Ecol. Notes. 7:544–548.

Jombart T. 2008. Adegenet: A R package for the multivariate analysis of genetic markers. Bioinformatics. 24:1403–1405.

Larkin M.A., Blackshields G., Brown N.P., Chenna R., Mcgettigan P.A., McWilliam H., Valentin F., Wallace I.M., Wilm A., Lopez R., Thompson J.D., Gibson T.J., Higgins D.G. 2007. Clustal W and Clustal X version 2.0. Bioinformatics. 23:2947–2948.

Lê S., Josse J., Husson F. 2008. FactoMineR: An R Package for Multivariate Analysis. J. Stat. Soft. 25:1–18.

Leigh J.W., Bryant D. 2015. POPART: Full-feature software for haplotype network construction. Methods Ecol. Evol. 6:1110–1116.

Li H. 2013. Aligning sequence reads, clone sequences and assembly contigs with BWA-MEM. arXiv. 1303.3997v2:1–3.

Lischer H.E.L., Excoffier L. 2012. PGDSpider: an automated data conversion tool for connecting population genetics and genomics programs. Bioinformatics. 28:298–299.

Liu X., Fu Y.-X. 2020. Stairway Plot 2: demographic history inference with folded SNP frequency spectra. Genome Biol. 21:280.

Marr J.C., Schaefer F.B. 1949. Definitions of body dimensions used in describing tunas. Fish. Bull. Fish. Wild. Serv. 51:241–244.

Meier J.I., Sousa V.C., Marques D.A., Selz O.M., Wagner C.E., Excoffier L., Seehausen O. 2017. Demographic modelling with whole-genome data reveals parallel origin of similar *Pundamilia* cichlid species after hybridization. Mol. Ecol. 26:123–141.

Muto N., Alama U.B., Hata H., Guzman A.M.T., Cruz R., Gaje A., Traifalgar R.F.M., Kakioka R., Takeshima H., Motomura H., Muto F., Babaran R.P., Ishikawa S. 2016. Genetic and morphological differences among the three species of the genus *Rastrelliger* (Perciformes: Scombridae). Ichthyol. Res. 63:275–287.

Noguerales V., Ortego J. 2022. Genomic evidence of speciation by fusion in a recent radiation of grasshoppers. Evolution. 76:2618–2633.

Patterson N., Moorjani P., Luo Y., Mallick S., Rohland N., Zhan Y., Genschoreck T., Webster T., Reich D. 2012. Ancient admixture in human history. Genetics. 192:1065–1093.

Petr M., Vernot B., Kelso J. 2019. admixr—R package for reproducible analyses using ADMIXTOOLS. Bioinformatics. 35:3194–3195.

Pickett B.D., Glass J.R., Johnson T.P., Ridge P.G., Kauwe J.S.K. 2022. The genome of a giant (trevally): *Caranx ignobilis*. Gigabyte. 2022:1–16.

Pritchard J.K., Stephens M., Donnelly P. 2000. Inference of population structure using multilocus genotype data. Genetics. 155:945–959.

R Core Team. 2022. R: A language and environment for statistical computing. https://www.r-project.org/.

Rambaut A., Drummond A.J., Xie D., Baele G., Suchard M.A. 2018. Posterior summarization in Bayesian phylogenetics using Tracer 1.7. Syst. Biol. 67:901–904.

Ruffley M., Smith M.L., Espíndola A., Turck D.F., Mitchell N., Carstens B., Sullivan J., Tank D.C. 2022. Genomic evidence of an ancient inland temperate rainforest in the Pacific Northwest of North America. Mol. Ecol. 31:2985–3001.

Stange M., Sánchez-Villagra M.R., Salzburger W., Matschiner M. 2018. Bayesian divergence-time estimation with genome-wide single-nucleotide polymorphism data of sea catfishes (Ariidae) supports Miocene closure of the Panamanian Isthmus. Syst. Biol. 67:681–699.

Su Y.-C., Su S.-H., Li H.-Y., Wang H.-Y., Lee S.-C. 2020. Implication of single year seasonal sampling to genetic diversity fluctuation that coordinates with oceanographic dynamics in torpedo scads near Taiwan. Sci. Rep. 10:16829.

Suyama Y., Hirota S.K., Matsuo A., Tsunamoto Y., Mitsuyuki C., Shimura A., Okano K. 2022. Complementary combination of multiplex high-throughput DNA sequencing for molecular phylogeny. Ecol. Res. 37:171–181.

Suyama Y., Matsuki Y. 2015. MIG-seq: an effective PCR-based method for genome-wide single-nucleotide polymorphism genotyping using the next-generation sequencing platform. Sci. Rep. 5:16963.

Tine M., Kuhl H., Gagnaire P.-A., Louro B., Desmarais E., Martins R.S.T., Hecht J., Knaust F., Belkhir K., Klages S., Dieterich R., Stueber K., Piferrer F., Guinand B., Bierne N., Volckaert F.A.M., Bargelloni L., Power D.M., Bonhomme F., Canario A.V.M., Reinhardt R. 2014. European sea bass genome and its variation provide insights into adaptation to euryhalinity and speciation. Nat. Commun. 5:1–10.

Wright M.N., Ziegler A. 2017. ranger: a fast implementation of Random Forests for high dimensional data in C++ and R. J. Stat. Soft. 77:1–17.

[dataset]* Santos S.R. 2014. Data from: “Complete mitochondrial and partial nuclear genomes for the jack species *Caranx ignobilis* (Forsskål, 1775) and *C. melampygus* (Cuvier, 1833) (Perciformes: Carangidae) from the High Hawaiian Islands” in Genomic Resources Notes accepted 1 October 2013 – 30 November 2013. Dryad. https://doi.org/10.5061/dryad.6gr7t.

**Appendix 2: Taxonomic implications**

The discovery of distinct morphological characters would allow us to describe the three clusters as taxonomically distinct species. Tentatively, the West cluster is diagnosed by the number of gill rakers (less than 29 in West vs. more than 31 in East and North) (Fig. 3f). The East and North clusters are separable primarily by the distances between the tip of snout and the 1st and 2nd dorsal fins (1st: 31.5–33.3% of standard length in East vs. 33.1–35.7% in North; 2nd: 46.2–48.3% in East vs. 47.8–50.3% in North).

*Megalaspis cordyla* was originally described as *Scomber cordyla* by Linnaeus (1758) from an unknown locality, with an erroneous note that the species was distributed in "America". Four nominal species are currently considered junior synonyms of *M. cordyla*: *Scomber rottleri* Bloch, 1793 (type locality: East-India); *Scomber guala* Lacepède, 1800 (South America) (unnecessary replacement name of *S. cordyla*); *Caranx besselii* Agassiz, 1874 (Indian Ocean); *Citua plumbea* Quoy & Gaimard, 1825 (Mauritius) (Fricke et al, 2024). Due to the absence of type specimens of some nominal species and the lack of detailed morphological information in the original descriptions (Kottelat , 2013; Fricke et al., 2024), applicable names for the three clusters are unclear. However, the type localities provide more information. *Scomber rottleri*, originally described from “East India” (Indo-Malay Archipelago), could refer to both the East and West clusters (Fig. 1a). Similarly, *Ca. besselii* and *Ci. plumbea*, described from the Indian Ocean, could refer to the East and West clusters, given that their distributions possibly extend to the eastern part of the Indian Ocean (Muto et al., unpublished data). Contrastingly, these three names could not be associated with the North cluster due to its restricted distribution range. Finally, *M. cordyla*, with an undetermined original locality, could be associated with any of the three clusters. Further taxonomic considerations are needed to determine valid scientific names for the three clusters.

*References*

Agassiz L. 1874. Pisces Celebes. Icones Piscium. London. Pls. 1–2, 4–5.

Bloch M.E. 1793. Naturgeschichte der ausländischen Fische. Berlin. v. 7: i–xiv + 1–144, Pls. 325–360.

Fricke R., Eschmeyer W.N., Van der Laan R. (eds) 2024. Eschmeyer's Catalog of Fishes: genera, species, references. http://researcharchive.calacademy.org/research/ichthyology/catalog/fishcatmain.asp

Kottelat M. 2013. The fishes of the inland waters of southeast Asia: a catalogue and core bibliography of the fishes known to occur in freshwaters, mangroves and estuaries. Raffles Bull. Zool. Suppl. 27: 1–663.

Lacepède B.G.E. 1800. Histoire naturelle des poissons. v. 2: i–lxiv + 1–632, Pls. 1–20.

Linnaeus C. 1758. Systema Naturae, Ed. X. (Systema naturae per regna tria naturae, secundum classes, ordines, genera, species, cum characteribus, differentiis, synonymis, locis. Tomus I. Editio decima, reformata.) Holmiae. v. 1: i–ii + 1–824.

Quoy J.R.C., Gaimard J.P. 1824–25. Description des Poissons. Chapter IX. In: Freycinet, L. de, Voyage autour du Monde...exécuté sur les corvettes de L. M. "L'Uranie" et "La Physicienne," pendant les années 1817, 1818, 1819 et 1820. Paris. 192–401 [1–328 in 1824; 329–616 in 1825], Atlas pls. 43–65.
